# Hierarchical motion perception as causal inference

**DOI:** 10.1101/2023.11.18.567582

**Authors:** Sabyasachi Shivkumar, Gregory C. DeAngelis, Ralf M. Haefner

## Abstract

Since motion can only be defined relative to a reference frame, which reference frame guides perception? A century of psychophysical studies has produced conflicting evidence: retinotopic, egocentric, world-centric, or even object-centric. We introduce a hierarchical Bayesian model mapping retinal velocities to perceived velocities. Our model mirrors the structure in the world, in which visual elements move within causally connected reference frames. Friction renders velocities in these reference frames mostly stationary, formalized by an additional delta component (at zero) in the prior. Inverting this model automatically segments visual inputs into groups, groups into supergroups, etc. and “perceives” motion in the appropriate reference frame. Critical model predictions are supported by two new experiments, and fitting our model to the data allows us to infer the subjective set of reference frames used by individual observers. Our model provides a quantitative normative justification for key Gestalt principles providing inspiration for building better models of visual processing in general.

## 1 Introduction

If motion can only be defined relative to a reference frame (Hoefer et al., 2023), what is the brain’s reference frame for the perception of a moving object? A century of psychophysical studies has provided us with evidence that motion is alternatively perceived in a retinotopic reference frame (Adelson and Bergen, 1985), in allocentric (world) coordinates (Warren and Rushton, 2009), or coordinate frames defined by other objects in a visual scene (Johansson, 1950; Restle, 1979; Gogel and Koslow, 1972; Gershman et al., 2016; Wu and Spering, 2022). Interestingly, perceived motion can rarely be explained by a single reference frame. For instance, in the famous Johansson illusion (Johansson, 1950), while the perceived velocity of the center dot is clearly biased away from the observed retinal velocity, it is not vertical as predicted by a reference frame defined by the flanker dots. Equally, the “flow-parsing” hypothesis (Warren and Rushton, 2009) proposes that the brain subtracts optic flow signals that are compatible with self-motion in order to make us perceive object motion in allocentric coordinates; yet, the empirically observed subtraction is rarely complete (Niehorster and Li, 2017)

Importantly, our perception of motion appears to be closely linked to an observer’s perception of the “Gestalt” of a scene, its structure, or configuration (Wertheimer, 1912). Recent work has made some progress in mathematically formalizing this elusive concept of a Gestalt: first in information-theoretic terms (Pomerantz and Kubovy, 1986; Restle, 1979), and more recently in closely-related Bayesian terms (Froyen et al., 2015; Feldman, 2009; Jäkel et al., 2016). A Bayesian formulation, compatible with the widely influential idea that the brain combines incoming sensory information with prior expectations to form subjective beliefs about the outside world (Von Helmholtz, 1867; Knill and Richards, 1996), has the advantage that these priors can be justified by the statistics and structure of the outside world. Yet, existing Bayesian accounts of motion perception are either formulated in a purely retinotopic reference frame (Weiss et al., 2002; Stocker and Simoncelli, 2006) or use priors that are not justified by our knowledge about the world but instead by computational tractability (Jäkel et al., 2016). Furthermore, quantitative empirical tests of some models in the motion domain are based on explicit questions about structure identity like “Do you perceive structure A or B?” (Gershman et al., 2016; Yang et al., 2021) which are known to be susceptible to decision biases which need to be estimated separately (Noel et al., 2022). These biases could arise for example from categories presented according to a different distribution than observers’ beliefs. Other studies (Bill et al., 2020) employ a motion prediction task with reports in the retinotopic reference frame. This design avoids the problem of decision biases by inferring the structure from reports about a different question. However, while successful in identifying the structure, this design does not address how the motion is perceived.

On a mechanistic level, motion signals are processed locally in early visual areas, and the brain needs to combine information across these local motion detectors to form a coherent percept. A long line of research has modeled how these early areas detect local motion (Hassenstein and Reichardt, 1956; Adelson and Bergen, 1985) and how, through a series of linear-nonlinear stages, this local motion can be combined into a more global motion percept (Adelson and Bergen, 1985; Rust et al., 2006). While integrating these local motion signals allows the brain to solve the ‘aperture’ problem (Braddick, 1993), it is not always useful for the brain to integrate information. In fact, local motion differences are a powerful segmentation cue, and several studies (Gogel and Koslow, 1972; Braddick, 1993) have shown how our brain contrasts local motions to perceive relative motion. It is however unclear how the brain decides between these two opposing operations, integration and segmentation.

A separate line of research (Körding et al., 2007) in multisensory integration has modeled how the brain solves a similar problem of deciding when to combine information across cues in a Bayesian framework (‘causal inference’). Given the general nature of this problem of deciding when to combine information, causal inference has been proposed to be a universal computational motif across the sensory cortex (Shams and Beierholm, 2022). In motion perception, causal inference has been used to successfully explain biases in estimating heading direction from both visual and vestibular cues (Acerbi et al., 2018; Dokka et al., 2019), but not the perception of moving objects.

Here we present a hierarchical causal inference model that overcomes all of the above challenges by performing joint inference over the hierarchical structure of a scene and the motion of individual visual elements within it. Importantly, the motion priors in our model are justified by motion in the real world, in which most objects are not merely slow, but exactly stationary with respect to their canonical reference frame (Knill and Richards, 1996). We also present new data from two psychophysical experiments in humans that probe the hierarchical perception of motion and that provide strong support for the key elements of our model, in particular its hierarchical structure, novel prior, and approximate computations.

## 2 Results

### Motion is perceived in dynamically inferred reference frames that reflect the causal structure of the world

Much of the world consists of approximately rigid objects that in turn are made up of approximately rigid parts. During translation, all points on a rigid object move in the world at the same speed. Consequently, a common velocity for multiple moving elements in a visual scene acts as a strong cue that the elements belong to a single object. Not surprisingly, when we observe a group of dots moving at the same speed (Figure 1A, top), our brain combines them into one object that it perceives as moving (Figure 1A, bottom), rather than perceiving the individual parts as moving independently (Wertheimer, 1912; Jansson and Johansson, 1973). This common velocity cue also allows the brain to segment the scene into multiple moving objects. For example, when we observe the dots moving as shown in Figure 1B, top, we perceive two partially overlapping objects moving at their respective velocities (Figure 1B, bottom).

**Figure 1.**
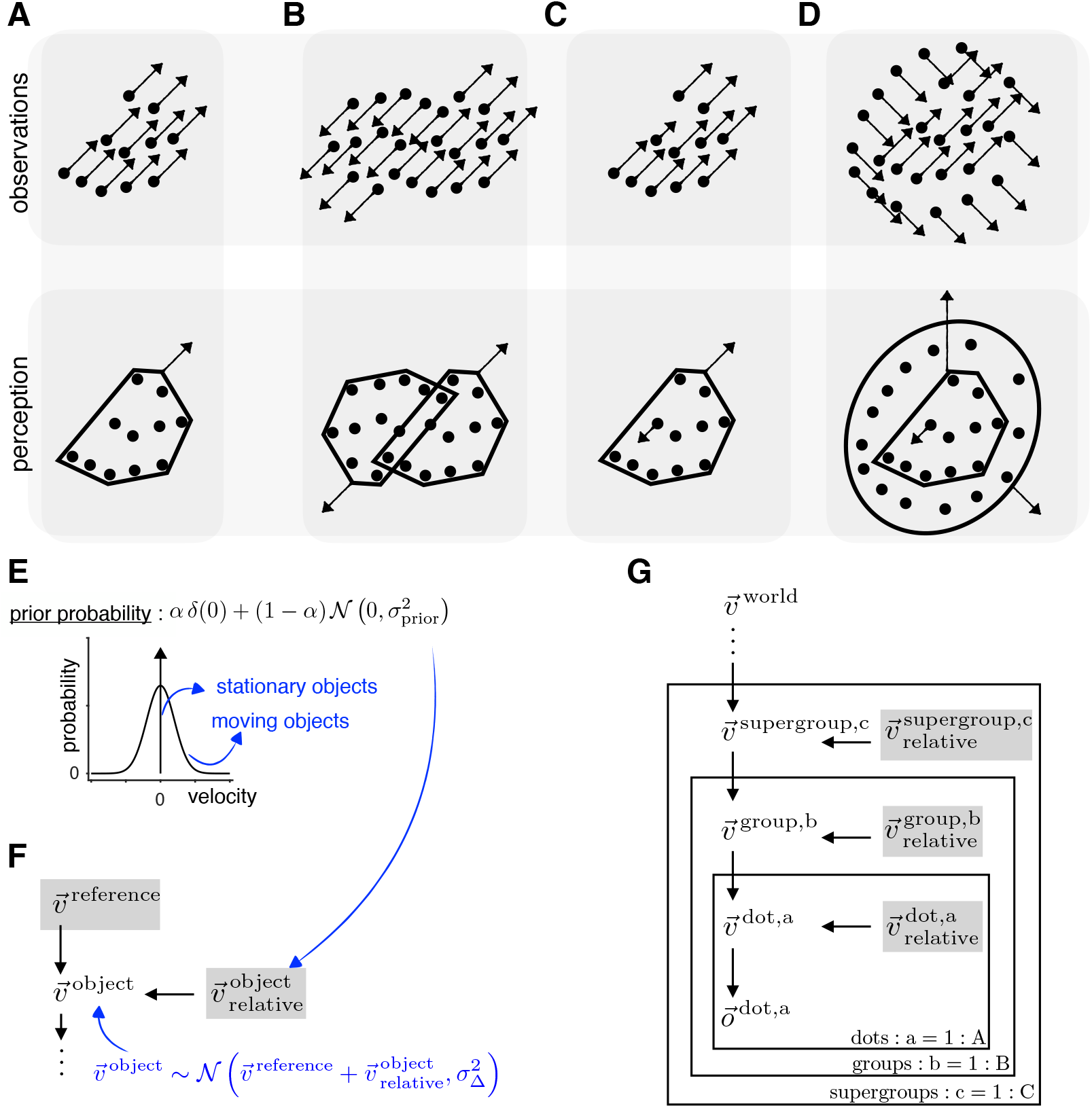
Causal inference model for hierarchical motion perception. **(A-D)** Illustration of four different dot patterns that are observed to move as shown in the top row with the arrows indicating retinal velocity vectors. The bottom row shows our predicted motion percept where the brain uses common velocity information to combine dots into objects and group objects into a hierarchical structure. **(E)** Prior that consists of a mixture of a delta distribution at 0 and a Gaussian distribution centered at zero, reflecting the knowledge that elements are either exactly stationary in an appropriate reference frame or are likely to move with a slow speed. **(F)** Generative model motif in which the object’s observed retinal velocity is the sum of the reference frame velocity 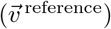 and its velocity with respect to the reference frame 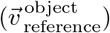. **(G)** Hierarchical causal inference model obtained by repeatedly applying the motif in F. Inference in this model leads to hierarchical grouping of dots, and representing dot motion in reference frames defined by the groups they belong to. The percept is determined by the non-zero relative variable lowest in the hierarchy.

Importantly, objects do not simply move independently in the world, but are related to each other through hierarchical whole-part relationships. When a part moves differently from the whole, the whole becomes a natural frame of reference in which to represent that part’s motion (Figure 1C-D). For example, the body is the natural frame of reference for the motion of an arm because of the causal whole-part relationship between the two: any change in the motion of the body is directly translated into a change in the motion of the arm.

Our key idea linking retinal observations to percepts in Figure 1A-D is that the brain dynamically constructs reference frames within which most of the visual elements it contains are stationary. We formalize this aspect of the physical world by extending the traditional slow-speed prior (Stocker and Simoncelli, 2006) over moving objects to include a mixture component consisting of a delta at 0. This is shown graphically in

Figure 1E, where ε denotes the prior probability that an object is stationary. This prior acts on the relative velocities of the visual elements represented in this reference frame. This brings us to the central motif in the generative model we propose is used by the brain to perform inference (Figure 1F). The motif specifies how the velocity of an object is the sum of the velocity of the reference frame and the velocity of the object within that reference frame. This sum is probabilistic, allowing for computational imprecision as quantified by 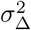. Inference within this generative model motif leads to a decomposition of the observed velocities of visual elements into the velocity of a (shared) reference frame and each element’s velocity relative to that reference frame. The degeneracy of this decomposition is broken by the mixture prior over the relative velocities, which leads to an automatic “chunking” of moving elements into groups that are inferred to move together. We hypothesize that the perceived velocity of a visual element moving in a reference frame is its relative velocity to the reference frame velocity if the relative velocity is non-zero and the reference frame velocity otherwise. This is compactly illustrated in our model by adding a shaded gray box around the candidate variables for perception in the generative model. Under this illustration, the percept corresponds to the candidate velocity that is non-zero and lowest in the model hierarchy (closest to the observations).

Recursively applying this motif leads to our proposed hierarchical causal inference model describing velocity percepts in a scene consisting of dots moving according to hierarchical causal relationships. Our model combines dots into groups, groups into supergroups, and so on (Figure 1G). We define a grouping tree as the graphical tree in which nodes at each level are the groups and supergroups inferred by the model, and the edges show which dots form groups, which groups form supergroups etc. The hierarchical causal structures relating the velocities in the scene are defined by specifying which nodes in the grouping trees are stationary (forming reference frames) and which nodes move in the appropriate reference frames (see Supplementary Information S2). At the bottom of the model are the actually observed velocities in retinal coordinates, 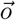, which are linked to the latent variables 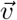 by a Gaussian likelihood whose width represents the observational noise. The top level of the hierarchy is the velocity corresponding to the stationary objects in the world in the egocentric reference frame. For stationary observers this is zero, but for moving observers it is equal to the optic flow velocity caused by self-motion. This allows for a natural extension of this model to explain deviations in perceived velocities due to self-motion (Warren and Rushton, 2009).

Performing Bayesian inference in this generative model requires computing a posterior belief over all possible causal structures in which the visual elements could be grouped, and over all the velocities in each of the structures. Before empirically testing the quantitative predictions of this model, we next explain the intuitive impact of its key elements using an increasing number of moving dots, building up to explain the classic Johansson illusion (Supplementary Video 1).

### Illustrating the causal inference model for dot stimuli consisting of 1-3 dots

We illustrate the key features of the model by applying it to very simple stimuli consisting of two or three dots. The model infers full posteriors over all possible structures and the velocities within each structure. For compactness, we focus on the most probable structures and show the most likely inferred velocity using vectors instead of variables in the generative model, as shown in Figure 2A for a simple stimulus in which a single dot is moving. We explicitly show the variables for the rest of the structures in Figure S1.

**Figure 2.**
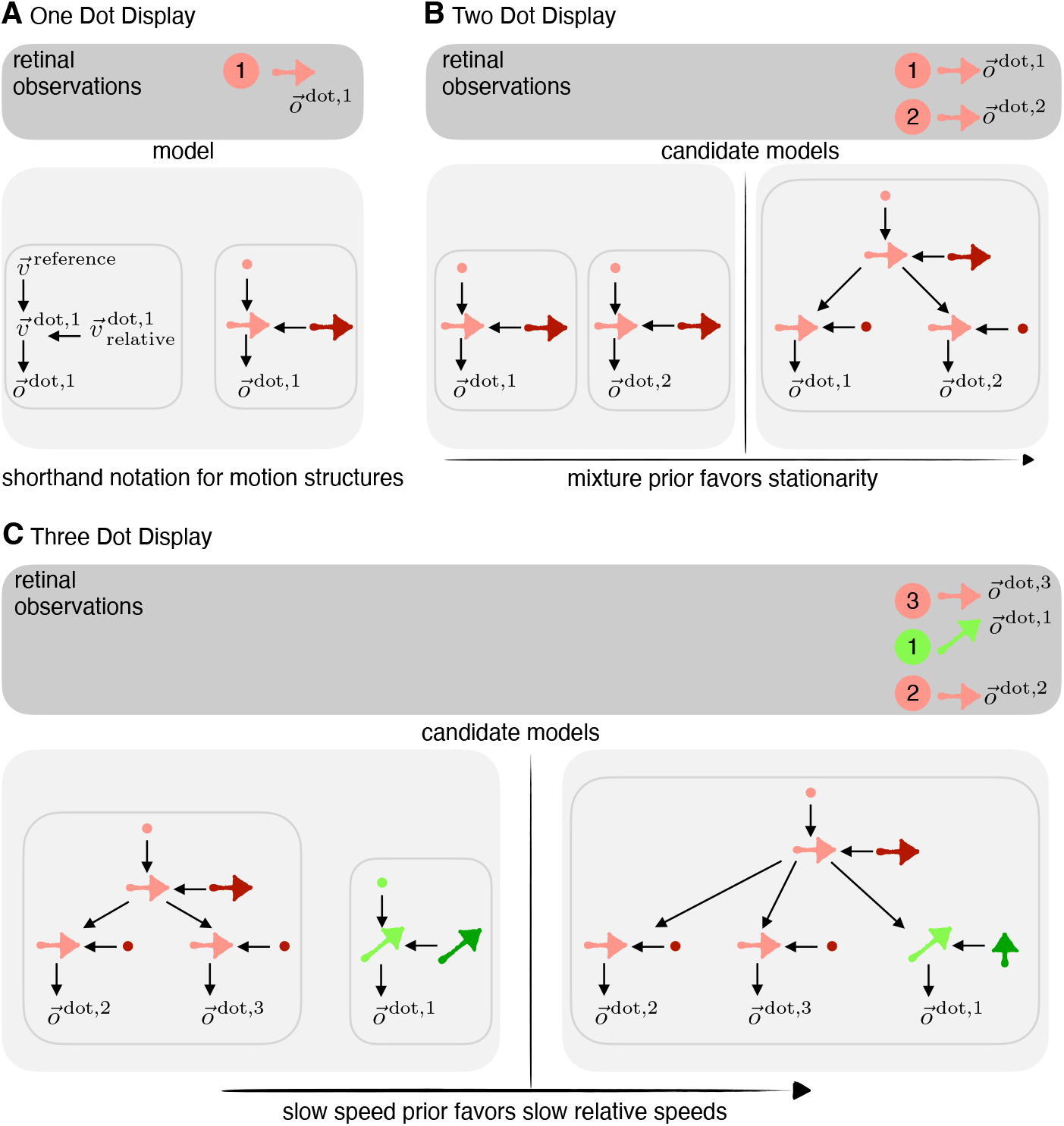
Model predictions for simple dot stimuli. **(A)** The motion of a single dot is inferred in the reference frame of a stationary world. In our shorthand notation, velocity variables have been replaced by their most likely value shown as a motion vector. Darker shades are used to indicate relative velocities and filled circles indicate zero velocity. **(B)** Two moving dots are explainable by two possible structures (left and right). If they move coherently, such that both are stationary with respect to a moving group, the delta component in the prior implies that most posterior mass lies on the combined structure (right). As a result we perceive a moving object consisting of both dots. **(C)** If a third dot is added to the display in (B), the observations are explainable by 264 different structures (Methods, and S2), two of which are shown here. On the left, the green dot is perceived as independent of, and unaffected by the motion of the red dots. On the right, the green dot is part of the same structure as the red dots, and perceived in a reference frame defined by a group in which two out of the three dots are stationary (favored by the delta component in the prior). The Gaussian component of the prior favors the right structure over the left one since the velocity of the green dot in the reference frame defined by the red dots is smaller than its velocity with respect to the stationary world. This explains the Johansson illusion (Johansson, 1950).

When observing two dots that move with the same velocity, there are two likely structures that can explain the observations: one in which both dots are moving independently, each represented in the egocentric coordinate system as an individual dot (Figure 2B, left structure), or a structure in which each dot can be inferred to be stationary with respect to a group (an abstract object), consisting of both dots, with the velocity of the group corresponding to the retinal velocity of the dots (Figure 2B, right structure). The delta component of our mixture prior over the relative velocities ensures that the latter hierarchical structure has the highest likelihood given the data since it has the fewest non-zero relative velocities. Furthermore, since the observed velocities for each dot will slightly deviate from each other due to observation noise, the group velocity combines both observations to obtain a more reliable velocity percept of the group (Trommershauser et al., 2011). This combination of dots into a single group occurs for all dots in the scene that are inferred to move with the same velocity.

If a third dot is added to the scene that moves with a different velocity from the other two dots, the two likely structures to explain the observations are: (a) an object consisting of the two coherently moving dots plus an independently moving third dot (Figure 2C, left) and (b) a single object consisting of all three dots in which the differently moving dot is represented as moving in the object’s reference frame (Figure 2C, right). The slow speed component in our mixture prior favors the latter structure if the differently moving dot has a smaller speed in the object’s reference frame than in the stationary egocentric reference frame. For instance, in the Johansson illusion (Figure 2C), the third dot has a small relative velocity with respect to the two coherently moving dots, and its perceived velocity is indeed biased towards its velocity in the reference frame provided by the group made up of the two dots (dark green vertical arrow in Figure 2C, right). However, it is important to recognize that even for as little as three moving elements, there are 264 different causal structures, for instance one in which dots 2 & 3 move relative to dot 1 rather than the other way around, or where all dots move relative to a reference frame defined by all of the dots together, moving at an intermediate velocity (for more information, and a derivation of the number of causal structures for n moving elements, see Supplementary Information S2). So it may not be surprising that structure (b) alone cannot explain human observers’ percepts which sometimes correspond to the retinal velocity suggesting structure (a), sometimes correspond to the relative velocity (structure (b)), and sometimes lie in between (Hochberg and Fallon, 1976; Shum and Wolford, 1983), suggesting that the brain performs inference over multiple, if not all, of these possible structures.

Furthermore, model predictions will depend on how perception is related to the posterior over structures and velocities. Prior work (Gershman et al., 2016) has suggested the mean for the most likely structure, but it could also correspond to the mean across structures as in other work on causal inference (Körding et al., 2007), or posterior sampling (Vul et al., 2014; Wozny et al., 2010).

### New empirical tests of model predictions

In order to quantitatively distinguish between our model and previous proposals, we designed two motion estimation tasks that tested the key elements of our model: (i) the novel mixture prior over relative velocity with a delta at zero, and (ii) the linking hypothesis mapping the posterior distributions over velocities in the model to the distribution over perceived velocities. Importantly, by using a motion estimation task, we test both causal inference over reference frames, and the perception of motion within a reference frame.

#### Experiment 1: Test using stimuli with two potential levels of hierarchy

In order to test our model, and to constrain its parameters, Experiment 1 was designed to precisely measure human motion perception during the transition from integration to segmentation. Observers used a dial to report their perceived motion direction of a patch of green dots, surrounded by a variable number of patches of red dots (Figure 3A). Surround dots were either stationary, or moving horizontally (direction 0 degrees). The number of surround patches was randomly chosen every trial from {1, 2, 3, 5, 10}, while the center always consisted of a single patch of dots. Additionally, the retinal center direction (direction on the screen) was randomly chosen on each trial from the set {0, ±2.5, ±5, ±10, ±20, ±45}°. The center and surround had a common horizontal velocity (0°), such that the direction of the center’s velocity relative to the surround was 90° depending on the sign of the center direction (more details in Methods). As expected, reported directions lie along the identity line when the surround is stationary (Figure 3B).

**Figure 3.**
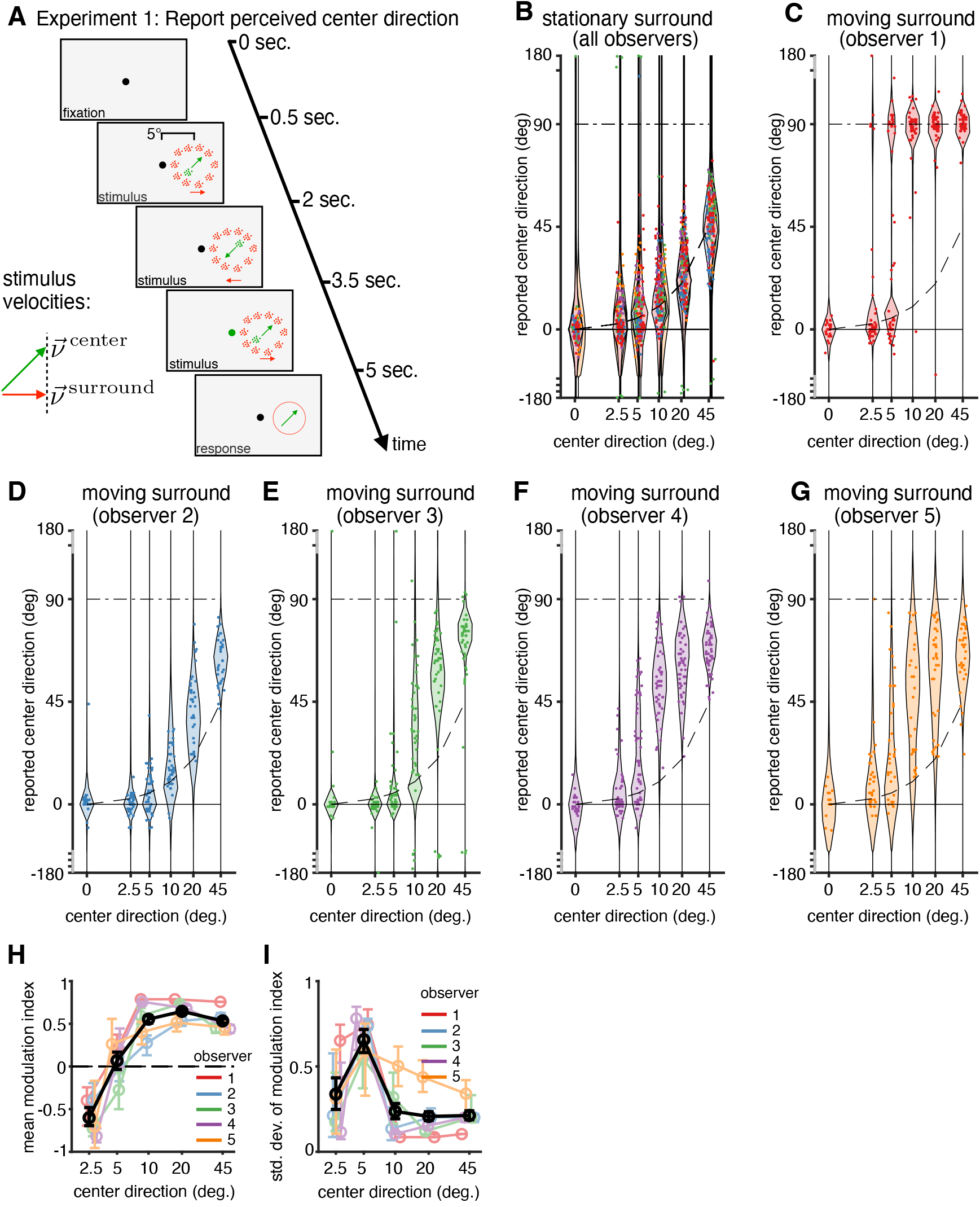
Experiment 1 – design and results. **(A)** During fixation, two groups of dots (red and green) appear and move back and forth three times for 4.5 seconds before disappearing. During the last phase of the movement, the fixation dot turned green. The observers adjusted a dial to report their perceived direction of the green dots during the last movement phase. The red (surround) dots were either stationary or moved horizontally (0°), while the green (center) dots varied in direction from trial to trial while keeping the horizontal component of their velocity matched with the surround. **(B)** Responses of all five observers overlaid for the condition where the surround is stationary. Each dot represents a single trial. All responses lie around the identity line (warped due to non-linear spacing of the x−axis). Responses were flipped for negative center directions to match the positive directions after verifying that the responses were symmetric. **(C-G)** Responses for each observer when the surround is moving. The horizontal lines at 0° and 90° indicate the predicted reports for complete integration (perceiving the surround) and complete segmentation, i.e., perceiving the relative velocity, respectively. **(B-G)** The overlaid violin plots show the model predictions (not data distributions). One model was fit jointly on all data for each observer. **(H**,**I)** Mean and standard deviation in modulation index (68% confidence intervals) defined such that −1 corresponds to pure integration, +1 to pure segmentation, and 0 to retinal motion. Different colors indicate different observers; black line denotes the average across observers.

When the surround is moving, observer responses systematically deviate from the identity line (Figures 3C-G, S4). Specifically, responses appear biased towards zero degrees (surround direction) for small center directions for most observers, consistent with the observer integrating the center and surround velocities and reporting the cue-combined velocity. The reported velocities are biased towards 90^°^ for larger center directions, consistent with the observer perceiving the relative velocity between the center and surround.

We quantify the effect of the surround motion on the center motion reports by mapping the responses to a modulation index. This index lies between −1 and +1, where −1 corresponds to complete integration (perceiving the surround direction), 0 corresponds to the surround having no effect, and +1 corresponds to complete segmentation (perceiving the relative velocity between the center and surround; see Methods for details). The mean modulation index, averaged across observers (Figure 3H) is negative indicating integration, for 2.5° (*p* < 0.001 for the group, *p* < 0.05 individually for 4 out of 5 observers, based on 5000 bootstrapped samples). For larger separations (greater than 10°), the average and individual mean modulation indices are positive indicating segmentation for larger separations (p < 0.001). The standard deviation of modulation index, averaged across observers (Figure 3F) is largest for intermediate separations (p < 0.01 for the difference in the standard deviations of the modulation index at 5° and at 2.5° and 45°) indicating a higher variability due to uncertainty over causal structures, in agreement with our model predictions.

Finally, we fit our model to the data by inferring posterior distributions over the model parameters given the data from each observer. We find that the empirical responses are in excellent agreement with the response distributions predicted by our causal inference model, which are overlaid as violin plots in Figures 3C-G. The goodness of fit was measured by variance explained (VE) to be between 92-96% across observers. Also note the clear evidence of causal inference – uncertainty whether to integrate or segment – in the form of bimodal responses (or increased variability) for intermediate center directions around 5° visible in both empirical responses and model fits. These characteristic deviations between retinal motion and perceptual reports depend on the number of surround patches, also in agreement with the model fits (Figure S3 with an average VE of 94%).

#### Insights from model fitting

We next fit the model to individual responses and obtained posterior distributions over the parameters for each observer (Methods). This allowed us to gain three key insights about the model: (a) whether the mass in the delta component was required to explain the pattern of responses, (b) how observers map the inferred posteriors to responses on each trial, and, (c) how the different causal structures contribute depending on center direction.

Remarkably, for all observers, most of the prior mass was in the delta-component, indicating the strength the brain’s expectation that relative motion in the world is exactly zero, rather than merely slow (Figure 4A). We further confirmed the presence of a delta component in the mixture prior by a formal model comparison (Figure 4B, leftmost column) and found strong evidence against a model without as compared to models with mass in the delta component.

**Figure 4.**
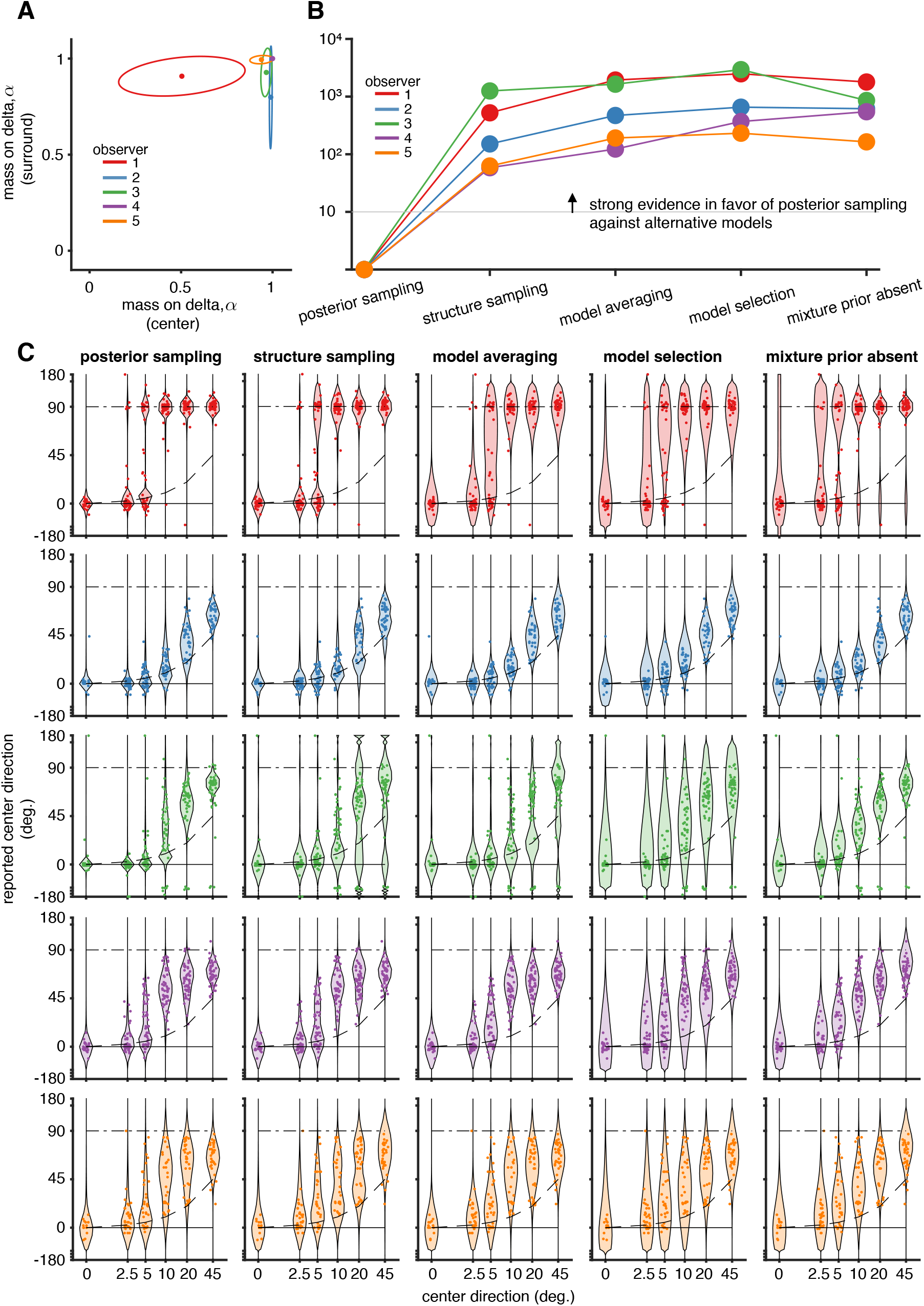
Model fitting insights from experiment 1. **(A)** Fitted mass on the delta component in the prior for center and surround. Each colored line show the mean and 95% CI for each observer. **(B)** Model comparison. For each model, the difference in AIC score to the posterior sampling model is shown. We find strong evidence against all alternative models as compared to the posterior sampling model. **(C)** Model predictions overlayed with data for each model considered in **B**.

Our model comparison (Figure 4B) also revealed that observers’ responses are best described as arising from approximate inference (posterior sampling) in the full Bayesian model. We compared this to other previously proposed maps from posteriors to responses: (a) reporting the posterior mean (model averaging) which is the optimal strategy for estimation tasks (Körding and Wolpert, 2004), (b) reporting the conditional mean under the most probable structure (model selection) which maximizes consistency (Stocker and Simoncelli, 2007) and minimizes the description length (Chater, 1996), and (c) reporting the mean by sampling the structure (structure sampling) which is a more precise approximation than posterior sampling (Wozny et al., 2010). Out of these models, previous studies on hierarchical motion perception (Gershman et al., 2016; Bill et al., 2022) have mapped posteriors to reports using the model selection strategy.

The predicted response for each of these models is shown in Fig. 4C overlaid with observer responses. Structure sampling reports the mean under the sampled structure. Thus any variability conditioned on the structure must be due to observation and motor noise which are shared across all structures (and cannot be ascribed to variability due to sampling conditioned on the structure, as for posterior sampling). This leads to larger-than-required predicted variability for some center directions, reducing the likelihood of the data under this model. Model averaging which reports the weighted mean across structures, increases this problem since averaging reduces variability. Furthermore, it also predicts intermediate reports which struggle to explain bimodal features in the data. Model selection reports the posterior mean within the structure with the highest posterior probability. While able to explain bimodal reports due to observation noise varying which structure is most likely, it overestimates observation and motor noise for the same reason that structure sampling and model averaging do: the low variability of the mean velocity conditioned on the structure. The mixture-prior-absent hypothesis is the same as “posterior sampling” but with no mass on the delta component in the mixture prior. Due to the absence of the delta prior, the predicted responses show weak segmentation, and increased observation noise in order to capture the increased variability in the data during the transition.

We also performed model comparison allowing for different prior parameters for different surround conditions and different elements in the stimuli (Figure S7). This allows us to account for influence due to other perceptual factors not accounted by our model. For all observers, the model favored by the evidence had different prior widths and different masses on the delta component for different velocity variables (center, surround, group), performed joint inference over structure and motion, and converted posteriors into responses by sampling.

#### Inferred structures in experiment 1

Our model also allowed us to determine the structures underlying each observer’s subjective percepts. We found that for all observers, only four out of 12 possible structures were assigned a significant posterior mass (Figure 5A-D for 10 surround patches, and Figures S5, S6 for all other conditions). Under structure 1, the observer integrates center and surround, thus perceiving the cue-combined velocity (Figure 5A). As expected, the posterior probability of this structure was highest when the center and surround moved with the same velocity and decreased with an increase in separation between center and surround velocities (Figure 5E). The bias in the perceived center velocity towards the surround was determined by the weight given to the surround (Figure 5I), which we quantified using the modulation index (as in the previous section). Under structure 2, center motion is perceived in the reference frame defined by the surround – the canonical structure typically assumed for center-surround motion segmentation (Figure 5B,F, J). Unexpectedly, this structure is dominant for only 1 out of 5 observers, plays a transient role for intermediate differences between center and surround motion directions for just 2 observers, and only plays a very minor role for the remaining observers. The same is true for structure 3, under which the surround is perceived in the reference frame of the center, and perception of center motion mostly coincides with retinal motion (Figure 5C,G,K). Finally, structure 4 implies that both center and surround motion are perceived with respect to a reference that moves at a velocity intermediate to both center and surround (Figure 5D,H,L). This is the structure that carries primary responsibility for intermediate percepts (i.e. the apparently incomplete subtraction of the surround from center velocity) at large differences between center and surround motion directions. Surprisingly, none of the observers places any mass on the possibility that center and surround might belong to different causal structures (the 99th percentile of mass on this structure is below 1% for all observers) – an alternative potential explanation for intermediate percepts. However, one prediction of such a structure would have been to find a substantial fraction of responses along the identity line (given that responses are best explained by posterior matching overall), something that we did not observe (Figure 3C-G). The reason that structure 4 has higher posterior mass than structures 2 and 3 for large separations is the Gaussian slow speed component in the prior: the smaller relative velocities under structure 4 overpower the mass in the delta component of the prior.

**Figure 5.**
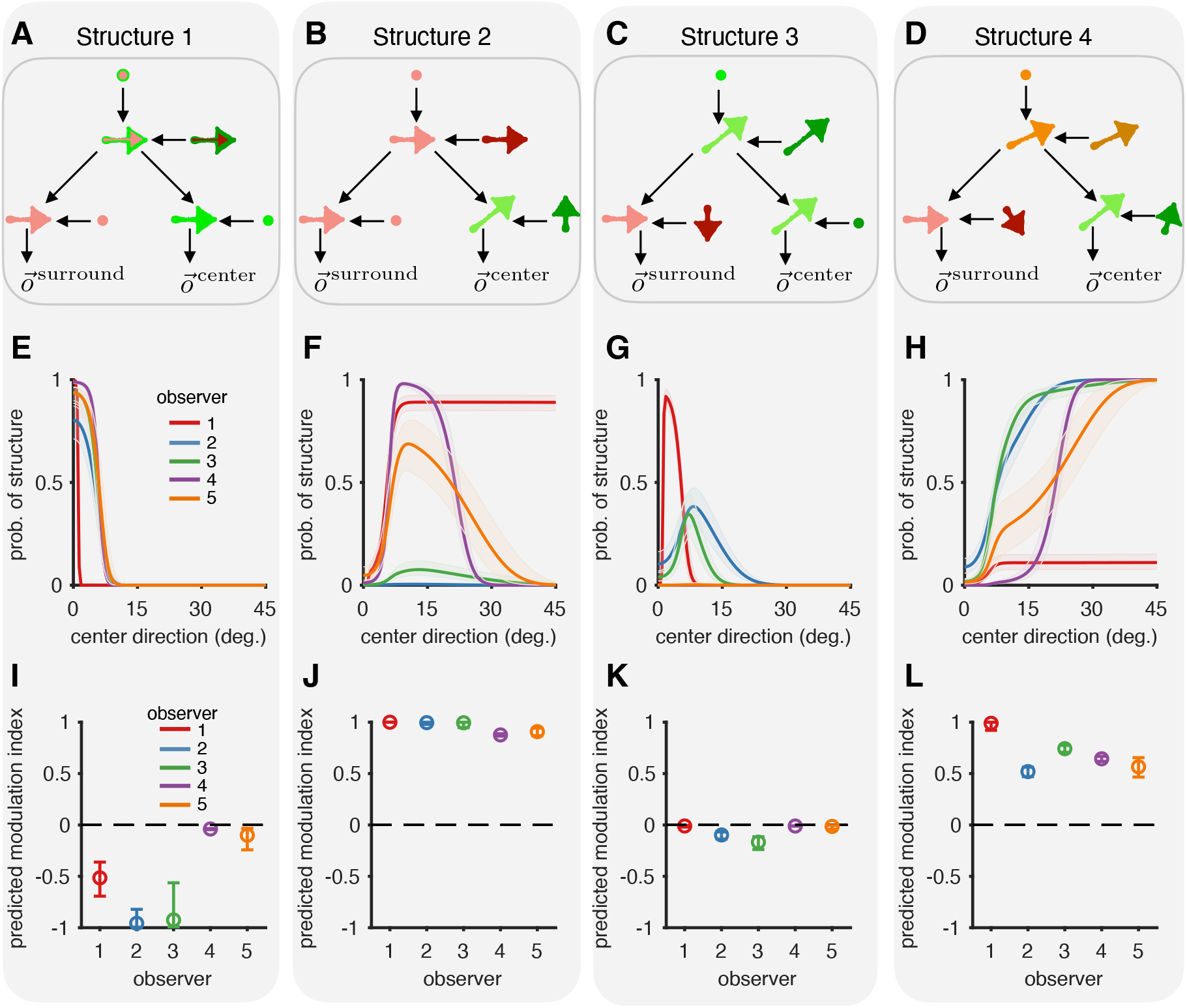
Inferred structures for observers in experiment 1. **(A-D)** The four causal structures that have a non-zero probability in our model fits. (See main text for description.) **(E-H)** The posterior probability (along with 95% CI) assigned to each structure by each of the observers as a function of center direction. All observers integrate the center and surround for center directions close to zero (C+G) and segments the center and surround otherwise. However, they differ to what degree they rely on each of the three different reference frames implied by D, E, and F. **(I-L)** As in Figure 3E, the modulation index (along with 95% CI) predicted under each structure quantifies the influence of the surround on the perceived center direction. The modulation index for a structure is independent of the probability assigned to that structure. Together, they determine the influence of the surround on the center.

As predicted by the model, we also find that both the integration effect (quantified by the model predicted modulation index at 0°) and segmentation effect (quantified by the model predicted modulation index at 45°) become stronger with increase in the number of surround patches (Figure S8B and S8C respectively).

#### Experiment 2: Test using stimuli with three potential levels of hierarchy

Next, we added a second surround of moving dots to test three key qualitative behavioral predictions of the hierarchical replication of our causal inference motif (different grouping trees shown in Fig.S21. First, motion perception during segmentation is local, i.e. the perceived motion of an element inferred to be moving relative to a group is independent of other motion in the visual scene. Second, if a visual element is integrated with a group, then it inherits the reference frame of that group. Third, the retinal velocity of a group, and not the velocity with which the group is perceived, determines the reference frame velocity for the elements that are part of that group (compare the alternative model in Figure S17 to Figure 1G).

As in Experiment 1, observers reported the direction of the center patch moving in the presence of a surround but in Experiment 2, the surround consisted of inner and outer rings of dots (Figure 6A). The inner and outer ring velocities were on opposite sides of the center patch velocity with the inner ring moving at {0°, −3°, −10°, −30°, −45°} counter-clockwise from the center. The outer ring moved at 60^°^ counter-clockwise from the center when the inner ring moved at 0^°^ and the outer ring’s velocity was adjusted to maintain a constant relative velocity between outer and inner rings as the direction of inner ring was varied. This design was motivated by the goal to find a clearer empirical signature of the integration process than possible in Experiment 1. When center and inner ring are integrated, we expect them to be perceived relative to the outer ring, leading to perceptual reports that are biased towards −90°. However, when the center is segmented from, and perceived relative to the inner ring, we expect perceptual reports biased towards +90°, just as in Experiment 1.

**Figure 6.**
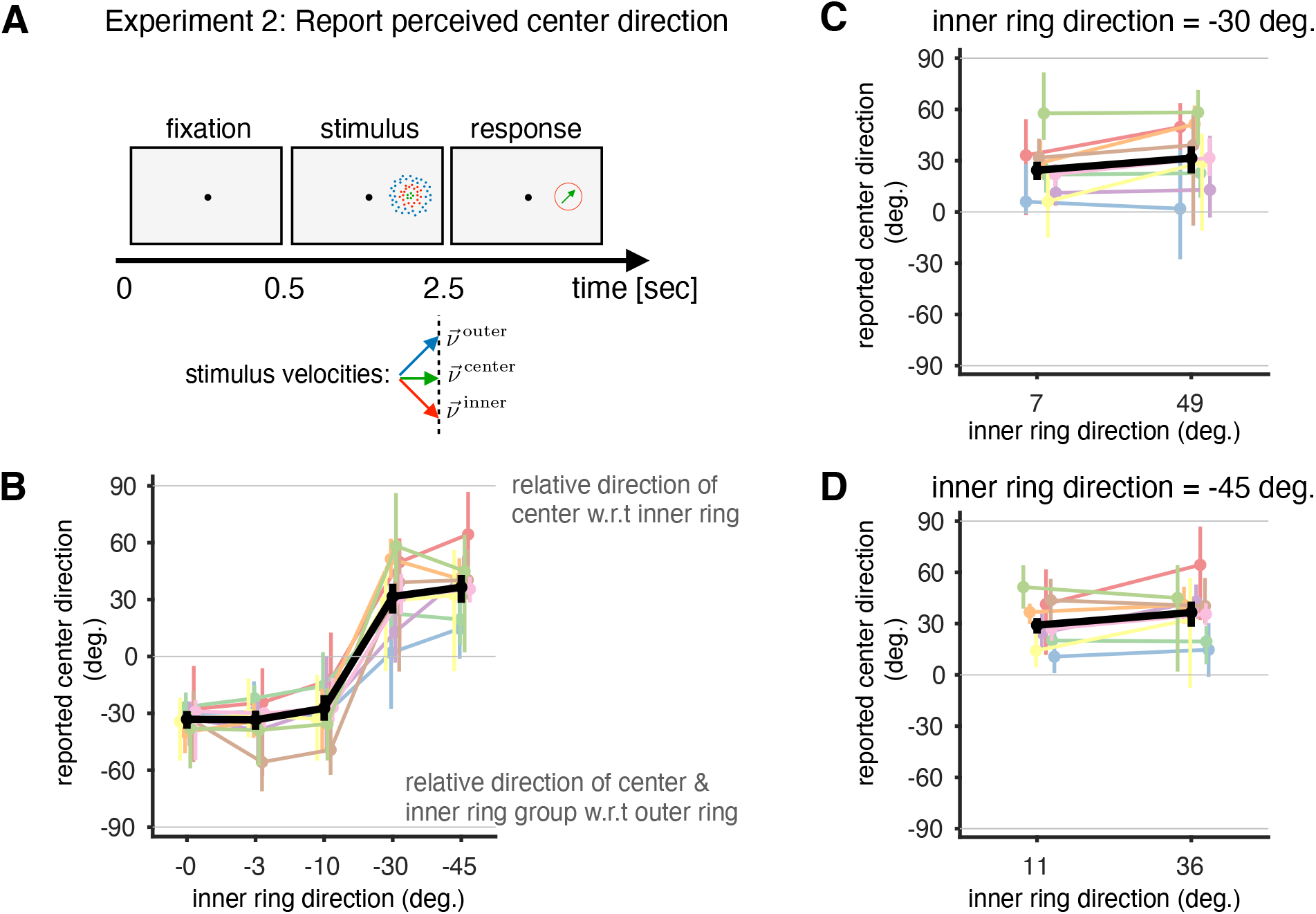
Design and results of Experiment 2 (three moving groups). **(A)** The observer performs an estimation task in which they have to report the direction of green dots (center) using a dial. While center dot directions are randomized from 0 to 360°, results are combined after rotating all velocities such that 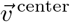 moves horizontally (0°). **(B)** Our model predicts that we will perceive the center dots in the inner ring’s reference frame if both move in noticeably different directions (segmentation, orange box) and cue combine the center and the inner ring motion if they are sufficiently similar (integration, magenta box). In the latter case, our percept would be the cue-combined center and inner ring’s velocity in the outer ring’s reference frame. The observer responses (circular median plotted with 95% CI) support these model predictions. **(C)** The causal inference model predicts that during segmentation, the percept of center motion should only depend on the retinal inner ring motion, not the perceived inner ring motion. The data (circular median plotted with 95% CI) clearly supports this prediction since the outer ring motion has no influence on the reported center directions (even though it influences the inner ring percepts, see Figure S18). **(D)** Same as (C) but for conditions where the inner ring moves at 45° and the outer ring moves at either 11° or 36°. We consider the two inner velocities that are most different from the center in our experiment to minimize the probability that the observer integrates the center and the inner ring velocities.

Since there are many more possible causal structures that could explain our stimulus, we would ideally make quantitative predictions using our full model. However, since we found this intractable due to large number of causal structures (264, see Supplementary section E), we generated qualitative predictions (Fig. S10) by making two simplifications: First, we only included structures in which every group had at least one stationary member (‘pure’ structures, Figs. S11-S16) – all the remaining structures predict percepts in between the percepts of these pure structures. Second, we assumed that the uncertainty about the inner ring velocity was much lower than that for the center, and that the uncertainty about the outer ring velocity was much lower than that about the inner ring – justified by the relative sizes of center, inner ring, and outer ring.

The data clearly confirmed our expectations, and qualitative model predictions: when the center and inner ring are integrated to form a group, this group is perceived in the reference frame formed with the outer ring (Figure 6B). In our stimulus design, the velocity of the group formed by the center and inner ring is always clockwise to the center ring velocity (with −90° the relative direction in the outer ring’s reference frame). We see clear signatures of integration of center and inner ring into a group for all observers (Figure 6B, p< 0.01) for smaller separations (≤3°). For larger separations between the center and inner ring, the model predicts that the center is perceived as moving in the reference frame formed with the inner ring, ignoring the outer ring. In our stimulus, this translates to reports being biased counter-clockwise to the center ring which we observer for the group (and 7/9 observers with p<0.01).

Our data also supports our model’s prediction that the retinal velocity of the inferred group determines the group’s reference frame velocity (Figures 6C and 6D respectively). Specifically, as the center is perceived relative to the inner ring when they move differently, the outer ring should have no effect on the percept. Consistent with this, we found no significant difference in reported center directions for different outer ring directions, for inner ring directions of 30° and 45° (p> 0.05 evaluated through bootstrapping with 1/9 observers showing significant difference for both inner ring directions). For these inner ring directions, the observers predominantly segment the center and inner rings (Figure 6B, orange box). The individual trial responses are shown in Figure S18.

In addition to the conditions shown in Figure 6B, we also test how: (a) the observers perceive the inner ring in the presence of the center and/or surround, (b) how the center ring percept is biased by just the inner ring moving coherently (similar to Experiment 1) and just the outer ring moving coherently. Responses in these conditions also agree with predictions of the causal inference model (Figure S19C-J).

## 3 Discussion

We have presented a new model of complex motion perception, and new data to support this model. The key normative ingredients, reflecting the structure of the physical world, are that the motion of visual elements should be represented in reference frames that they are causally connected to, that due to friction forces, the velocity in such a reference frame is mostly zero, and that the world is compositional and hierarchically organized. We designed two experiments in order to test all key elements of our model, to constrain its parameters, and to generate new insights for motion perception in scenes with rich motion structure.

Our work advances the large body of work trying to understand sensory processing in terms of natural input statistics (Attneave, 1954; Barlow and others, 1961; Hyvärinen et al., 2009; Laughlin, 1981). Whereas prior work started with the input statistics at the sensory periphery, our model explicitly incorporates causal relationships (moving the whole will move the part) and the importance of friction (the relative velocity is zero for most objects that are touching each other) for giving rise to these statistics. Strikingly, fitting our model to perceptual reports revealed that most prior mass is in fact at zero for all of our human observers (Figure 4A).

Our work was strongly inspired by Gershman et al. (2016) who embedded the vector decomposition idea of Johansson (1950) in a Bayesian inference framework and extended it hierarchically, and spatially. We changed their model in three ways: first, we changed the structure of the model in such a way that inference only required location computations. Second, this allowed us to define prior distributions over the relative velocities, and to use known physics to inform them by including a delta-component at zero. This prior over velocities also implied a prior over causal structures that differed from the prior based on the Chinese Restaurant Process assumed by Gershman et al. (2016) for mathematical convenience that places elements in a group in proportion to its size, or from related works that used a Jeffrey’s prior over motion strength to penalize the levels of hierarchy (Bill et al., 2022, 2020). Finally, we allowed for a flexible mapping from posteriors over structures and velocities to perceptual reports. Our data strongly constrained this mapping to correspond to sampling, different from the MAP inference over structures assumed by related prior works (Gershman et al., 2016; Bill et al., 2020; Yang et al., 2021; Bill et al., 2022).

The richness of our psychophysical data, together with formal model comparison, also allowed us to answer the important question of how the often-complex posterior distributions of a Bayesian observer are related to our deterministic-appearing percept (Block, 2018; Rahnev et al., 2021). In prior work, percepts were modeled as averages over velocity within the most probable structure only (‘model selection’, i.e. MAP inference over structures) (Gershman et al., 2016; Bill et al., 2022). Our experiments and analyses provide overwhelming evidence that, at least in the context of motion processing, the brain performs joint inference over both structure and motion, and that perceptual reports are best described by samples from this joint distribution (Figure 4B).

Approximate joint inference in our model can also explain why effect sizes in prior studies usually deviated from previously proposed theories like (Bayesian) Vector Analysis (Johansson, 1950; Shum and Wolford, 1983) or flow-parsing (Warren and Rushton, 2009): a range of different causal explanations are all consistent with the presented impoverished psychophysical stimuli. We confirmed a closely related prediction of our model – the increase in integration and segmentation effect sizes with the number of dots, i.e. the uncertainty in the surround (Zhou et al., 2020) of our experiment 1 (Figure S8).

The mixture prior over velocity that we propose extends the slow-speed prior previously proposed by Weiss et al. (2002) and quantitatively investigated by Stocker and Simoncelli (2006). Strikingly, we found that most of the prior mass was in the delta component not included in earlier work. Most likely, the delta component was not required to explain the data in Stocker and Simoncelli (2006) since the speeds used in their experiments were too large for the delta component to matter. On the other hand, their data suggested a prior shape of speeds with heavier tails than the prior used by us (our Gaussian prior over velocity implies a Raleigh distribution over speed). Future work may want to combine a delta at zero with a heavier-tailed distribution over velocity to better match data at large velocities.

Fitting our model to individual responses revealed unexpected variability in the causal structures inferred by different observers. We had expected that partial segmentation for large differences in motion direction between center and surround was best explained by observers’ lack of confidence that center and surround were indeed part of the same object. However, our quantitative analysis showed this not to be true: partial segmentation was actually explained by observers perceiving both center and surround to be slowly moving with respect to an abstract reference frame moving with an intermediate velocity. A corollary to this result is that the larger the integration bias (i.e. the more the reference frame velocity aligns with the surround), the larger is the segmentation bias (abstract reference frame velocity is the cue combined velocity of the center and surround), in agreement with earlier work (Wu and Spering, 2022). This result also demonstrated the importance of the ‘slow-speed’ component of the prior on relative motion – in contrast to the otherwise analogous spike and slab prior (Mitchell and Beauchamp, 1988) common in machine learning.

In our stimuli, center dots had a different color from the surround in order to minimize observer confusion about which motion direction to report. This difference in color might have increased the probability of segmenting the center from the surround. Using our experimental paradigm, and model-based analysis, the effect of color, as well as the effect of other cues like spatial relationships, disparity, etc. can be quantitatively studied, and related to model parameters like the prior probability of inferred the same motion, or the same causal structure. Furthermore, since our work was entirely based on the perception of motion directions, future work should also test our model predictions for speed perception.

While it has long been recognized that both integration and segmentation are key operations underlying motion perception (Braddick, 1993), our model shows that there are in fact two qualitatively different kinds of segmentation: (1) do two visual elements belong to the same causal structure, and, if the answer is yes, (2) is a visual element moving with respect to its reference frame? While our model answers both questions using the framework of ‘causal inference’ from multisensory integration (Körding et al., 2007), which has been proposed as a ‘universal computation’ for cortex (Shams and Beierholm, 2022), only the first question corresponds to a question about causality as statistically defined (Pearl, 2009).

Our model also lends itself to making predictions for neurophysiological data. We leave for future work the tantalizing possibility that the two kinds of variables in our model (corresponding to the left and right sides of Figure 1G) are represented by the two major classes of neurons who have been reported (Born and Bradley, 2005) in cortical motion area MT (surround-suppressed and not surround-suppressed).

Bayesian causal inference has recently been proposed as a unifying theory for neuroscience (Shams and Beierholm, 2022). Our model extends the simple ‘same’ or ‘different’ scenarios in previous causal inference work to hierarchical whole-part relationships reflecting the compositionality of the world (Lake et al., 2015). The computations at the lowest level of our model in fact resembles a recent probabilistic model of neural responses in primary visual cortex (Coen-Cagli et al., 2015), and the hierarchical architecture of our model directly suggests an equivalent one for the ventral stream, potentially allowing us to understand both dorsal and ventral stream as performing inference in closely related generative models.

## Methods

### Observers

Five naive observers participated in Experiment 1, and 10 naive observers participated in Experiment 2. Observers provided written informed consent, and were financially compensated for their time. Experiments were approved by the Office for Human Subject Protection (OHSP) at the University of Rochester (IRB number 0003909). 1/10 observers in Experiment 2 was excluded based on their large response variability (standard deviation greater than 30°) in the control condition and their data was omitted from further analysis.

### Experiment 1 details: two moving elements

The stimulus consisted of a ‘center patch’ of green dots, presented at 5 degree eccentricity. Dots were 0.1 degree in diameter and distributed uniformly within the patch with a density of 6.88 dots/degree^2^. The center patch had a radius of 0.68 degrees and was surrounded by a ring of radius 2.72 degrees, consisting of a variable number of patches of red dots (Figure 3A). Each surround patch had a radius of 0.54 degrees and a dot density of 10.91 dots/degree^2^. Dot displays were viewed binocularly, and no disparity cues were added, such thatall the dots moved in the plane of the display. The stimuli were presented on a 27-inch monitor with a refresh rate of 60 Hz and a resolution of 1920 1080 at a viewing distance of 105 cm. Eye movements were tracked using an Eyelink 1000 system and trials were discarded in which eyes moved within 1 degree of the center patch.

The stimuli were presented at an eccentricity of 5 degrees in the periphery. The number of dots in a patch was fixed to 10. There was one center patch and the number of surround patches was chosen in every trial from the set [1,2,3,5,10]. The center retinal direction was chosen from the set [0°, ±2.5°, ±5°, ±10°, ±20°, ±45°] and the surround was either stationary (stationary surround always had 5 patches) or moved at 0 degrees (horizontally rightwards) at a speed of 1 degree/sec.

After a fixation period of 0.5s, the stimulus appeared and moved back and forth for 1.5 cycles. The patch envelopes moved at a constant velocity and reversed their velocity after 1.5 seconds (square wave velocity profile with a time period of 3 seconds). The back-and-forth movement ensured that the envelopes stayed within a fixed area of the screen. In the last half cycle, the fixation dot turned green indicating that the observer had to report the direction during the last half cycle.

After stimulus offset, an arrow appeared at the location of the center patch, and observers used a dial to adjust the arrow direction to match their motion direction percept. The stationary surround condition served as a control. Observers who had a response standard deviation larger than 30 degrees in this condition were removed from subsequent analysis. The fixation period and stimulus had a total duration of 5 seconds following which the observers could make a response at any time to proceed to the next trial. Each observer participated in three sessions to get 22 trials per condition on average for a total of 1446 trials per observer on average.

### Experiment 2 details: three moving elements

In experiment 2, the stimulus consisted of a center patch, an inner ring, and an outer ring, all arranged concentrically centered at 5 degrees eccentricity (Figure 6A) separated by 0.5 degrees. On every trial, either the center dots, or the inner ring dots, were colored green, indicating whose direction had to be reported. The other parts of the stimulus contained red and blue dots, respectively, with the color assignment randomly drawn every trial. The number of dots in the center was 10 (density of 3.2 dots/degrees^2^ with a center patch radius of 1 degree), the inner ring contained 50 dots (density of 4 dots/degrees^2^ with a inner ring width of 1 degree) and the outer ring contained 250 dots (density of 11.4 dots/degrees^2^ with a outer ring width of 1 degree). The viewing distance, eye recording details, and the criteria for discarding trials due to fixation breaks and control condition response variability were the same as in experiment 1.

Each trial started with a 0.5 s fixation period during which only the fixation dot was shown on the screen. This was followed by the random-dot stimulus which moved for 2 seconds within an aperture at a constant velocity. Unlike for Experiment 1, the moving dots were presented inside fixed apertures, and the stimulus direction was not reversed. The center moved with a speed of 0.5 degrees/sec and the center direction was randomized across trials, within a range of 360^°^, to minimize the effect of reporting biases. The inner ring’s retinal direction was chosen from the set [0°, −3°, −10°, −30°, −45°] relative to the center where negative angles indicate clockwise rotations. The outer ring moved in a counter-clockwise direction chosen such that the same relative velocity was maintained between the outer and inner rings across different inner ring directions. Furthermore, the speeds of inner and outer ring were chosen such that the relative velocity between either ring and center was perpendicular to the center direction.

Different conditions were interleaved across trials in which the inner and outer rings could either move randomly or coherently and the observer had to report the direction of the center or inner ring. As in experiment 1, the observer made their responses by adjusting the direction of the arrow that appeared at the location of the center or inner ring after the stimulus offset with its size matched to the size of the corresponding target. Each observer performed three sessions to get 46 trials per condition on average for a total of 2239 trials per observer on average.

### Modulation index to summarize observer responses in Experiment 1

We defined the modulation index, w_MI_, such that the percept of the center patch 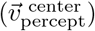 predicted for a given *w*MI is given by

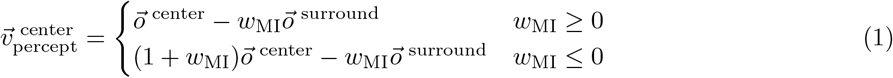

where 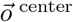 and 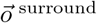 are the observed center and surround velocities from the brain’s perspective for a given trial. This definition incorporates partial subtraction of the surround (case w_MI_ ≥ 0) for relative velocities when the effect on perception is repulsive (Mori, 1979) and cue combination (Ernst and Banks, 2002) when the effect is attractive (case w_MI_ ≤ 0). Under the simplifying assumption that 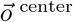 and 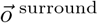 correspond to the experimenter-controlled velocities on the screen (ignoring observation noise), and that the perceptual reports exactly reflect the perceived variable 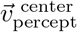 (ignoring motor noise), one could estimate a per-trial modulation index to obtain a distribution over w_MI_ for each separation of center and surround in order to estimate means and standard deviations which unfortunately would be biased. In order to obtain the unbiased estimates reported in Figures 3H and I, we therefore modeled observation noise, motor noise, and motor bias explicitly which allowed us to infer the distribution over w_MI_ from the distribution over perceptual reports (see supplementary section A for details).

### Causal inference model for hierarchical motion perception

#### Generative model in a scene with two moving elements

The observed retinal velocities, 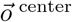 and 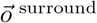, were modeled as the true retinal velocities, 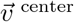 and 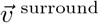 corrupted by additive Gaussian noise with variance 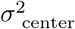 and 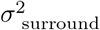, respectively (I refers to a 2 × 2 identity matrix) :

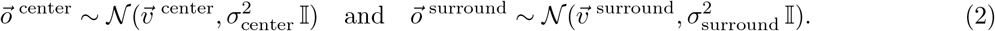

The velocities were parameterized as two dimensional vectors reflecting the x and y components. In order to model the inference over the different grouping trees (here, determining whether center and surround are part of the same moving group), we follow (Körding et al., 2007) in introducing a binary (logical) variable *S*^center,surround^ ∈ {0, 1} (corresponding to the left and right side of Figures 2B and S2B). This allows us to write the conditional probabilities compactly as:

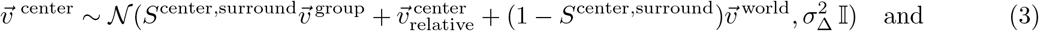

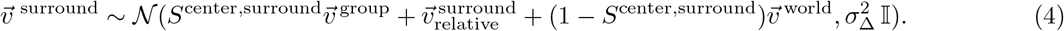

where 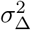 models the uncertainty in the velocity composition, e.g. due to computational noise. The prior over *S*^center,surround^ is given by *β*^center,surround^ which represents the prior probability that center and surround belong to a common structure (based on prior experience, or other non-motion cues). In a scene with only two moving elements, the group velocity is inferred in the stationary world reference frame with 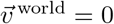 (Figure 1G):

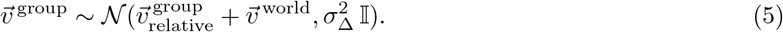

The prior over each of the relative velocities is a mixture prior of a delta function at zero and a normal distribution centered at zero

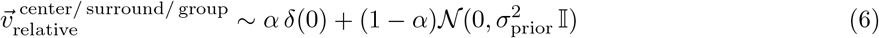

where ε represents the expectation that the (relative) motion is exactly zero (Figure 1E,F).

#### Mapping inferred latent variables to percepts

Our model predicts that if the center is inferred to be part of the same group as the surround, then the perceived center motion corresponds to the center’s relative velocity if it is inferred to be nonzero, and to the group velocity if the center’s relative velocity is inferred to be zero. If the center is not inferred to be part of the same group as the surround, then its motion relative to stationary world is perceived. By defining *C*^(*…*)^ ∈ *{*0, 1*}* to denote whether the velocity 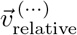 is zero (*C*^(*…*)^ = 0) or not (*C*^(*…*)^ = 1), we can compactly write the percept as:

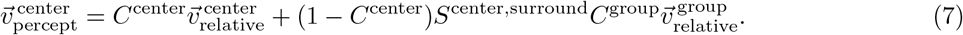

Therefore, the distribution over the observer’s percept is a mixture distribution with the mixture weights corresponding to the posterior probability of the different causal structures (characterized by possible *S*) and the different nested structures within each causal structure (characterized by possible *C*):

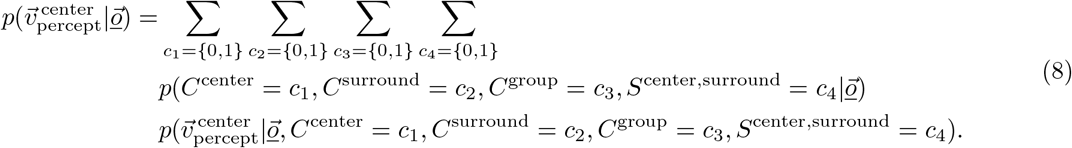

where for compactness we define 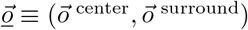. For any number of moving elements, this posterior has the general form:

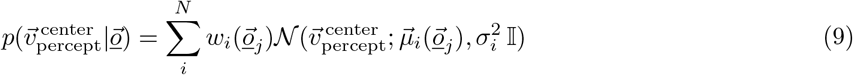

where *w*_*i*_, *μ*_*i*_, and 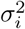 correspond to the probability, mean and variance associated with each of the *n* structures, respectively. For *n* moving elements, the total number of structures is bounded by 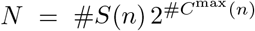 wherein #*S*(*n*) is the number of possible grouping trees, and 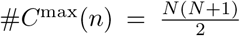 is the maximum number of nodes across all grouping trees. While this value of N overestimates the number of structures, the posterior unchanged from considering the exact number of structures (see supplementary section B for details). The expressions for the terms in Eq. 9 are derived in supplementary section C..

The distribution over perceived velocity can be mapped onto the distribution over the perceived center direction, 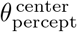, by mapping each velocity to its corresponding direction:

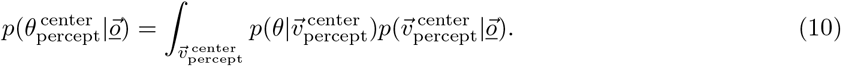

Inserting Eq. (9) into (10), we can approximate the integral by an analytic form (by approximating the distribution over directions implied by a Gaussian distribution over velocities by a von-Mises distribution) to get:

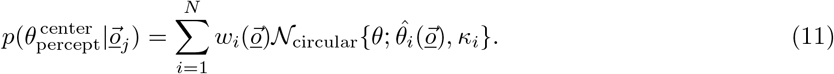

*N*_circular_ represents the von Mises distribution pdf with mean parameter 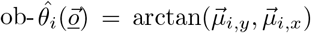 and concentration parameter 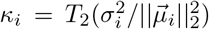. This approximation was obtained by numerically simulating two-dimensional velocities under Gaussian noise and approximating the corresponding distribution over directions using a von Mises distribution. We allow for lapses in responses by adding another component to the distribution over the center direction that is a von Mises distribution characterized by θ^lapse^ and κ^lapse^.

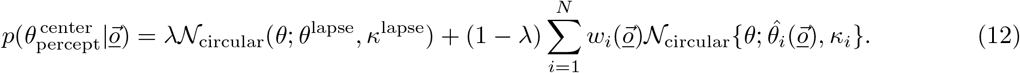

which can be simplified as

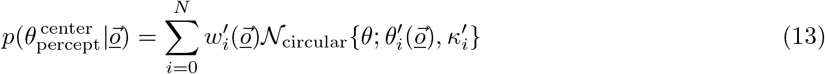

in which 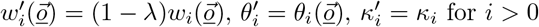, and 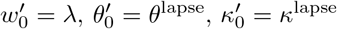.

#### Perceptual estimation

We consider the following four, previously proposed (Körding et al., 2007; Wozny et al., 2010), ways in which the brain may convert the posterior 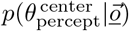 into a perceptual point estimate, 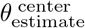:

**Model averaging:** 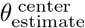 is the mean of the joint posterior over all structures: 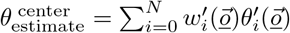.

**Model selection:** 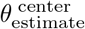 is the posterior mean over direction for the most likely structure: 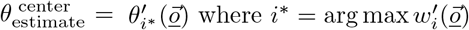

**Structure sampling:** 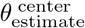 is the posterior mean over direction for a single structure sampled from the posterior over structures: 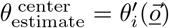 where probability of 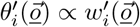.

**Posterior Sampling:** 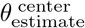 is a direction sampled from the joint posterior over all structures and directions: 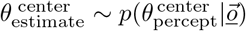.

#### Predicted distribution over observer responses

The distribution over observer reports, *R*, for a set of experimenter-defined directions 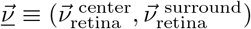 can be obtained by marginalizing out all possible sensory observations (Körding et al., 2007; Acerbi et al., 2014) :

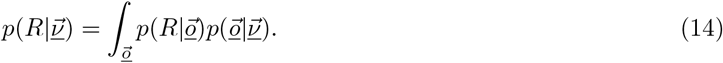

We model the distribution over observer reports, *R*, as a von Mises distribution centered on 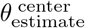 allowing for a reporting bias b and a motor noise κ_*m*_:

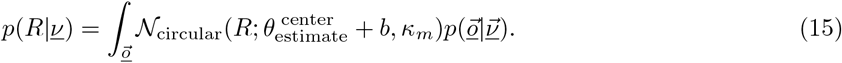

Since this integral is intractable to evaluate analytically, we approximate this using Gaussian quadratures evaluated at points 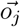 with weights 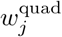:

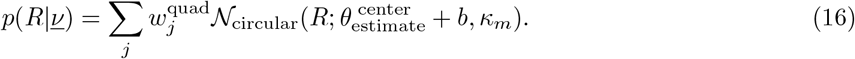

The distribution over observer responses for different perceptual estimates described in the previous section are given in Supplementary section D.

#### Model fitting details

We obtained the maximum a posteriori (MAP; for initializing the sampler) and maximum likelihood estimate (MLE; to compute AIC) for the model parameters under weakly informative priors (details in Table S1) using a quasi-newton Broyden-Fletcher-Goldfarb-Shanno (BFGS) unconstrained optimization procedure (fminunc in MATLAB). We obtained full posteriors over all model parameters using generalized elliptical slice sampling (Nishihara et al., 2014) which allowed us to get uncertainty estimates for all parameter estimates. We used 144 chains with 25000 samples per chain to estimate the posterior distribution over the parameters (average 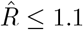).

To evaluate the absolute goodness of fit of the model, for each combination of center and surround velocities, we compared the empirical CDF from the reported directions with the corresponding model prediction. We quantified the overall match as variance explained across all conditions.

## Acknowledgements

We would like to thank Jan Drugowitsch, Gabor Lengyel, Boris Penaloza, Johannes Bill, Jean-Paul Noel, Greg Horwitz, Steven Grisafi, and Frank Jäkel for their helpful comments on our manuscript.

## Funding

This work was supported by:

National Institutes of Health grant U19NS118246 (to SS, GCD, and RMH)

Division of Information and Intelligent Systems IIS-2143440 (to RMH)

National Science Foundation grant NSF-1449828 (to SS).

## Author Contributions

Conceptualization: SS, GCD, RMH

Methodology: SS, GCD, RMH

Investigation: SS

Formal Analysis: SS, RMH

Visualization: SS, GCD, RMH

Funding acquisition: GCD, RMH

Writing – original draft: SS, GCD, RMH

Writing – review & editing: SS, GCD, RMH

## Code and Data availability

Code and data is available at https://osf.io/f9dsg/.

