## Supplementary Information for "Hierarchical motion perception as causal inference"

### Supplementary Text

#### A Estimating the distribution over modulation indices

In this section we present the details of the procedure to compute the distribution of modulation indices described in the main text. The predicted distribution over reports across trials for a given combination of veridical center and surround velocities can be expanded for a given modulation index $w_{\text{MI}}$ assuming Gaussian observation noise:

$$p(R|\vec{\nu}^{\text{ center}}, \vec{\nu}^{\text{ surround}}, w_{\text{MI}}) = \begin{cases} \mathcal{N}(R; b + \vec{\nu}^{\text{ center}} - w_{\text{MI}}\vec{\nu}^{\text{ surround}}, \\ \sigma^2_{\text{motor}} + \sigma^2_{\text{center}} + w^2_{\text{MI}}\sigma^2_{\text{surround}}) & w_{\text{MI}} \geq 0 \\ \mathcal{N}(R; b + (1 + w_{\text{MI}})\vec{\nu}^{\text{ center}} - w_{\text{MI}}\vec{\nu}^{\text{ surround}}, \\ \sigma^2_{\text{motor}} + (1 + w_{\text{MI}})^2\sigma^2_{\text{center}} + w^2_{\text{MI}}\sigma^2_{\text{surround}}) & w_{\text{MI}} \leq 0 \end{cases} \tag{S1}$$

with the corresponding CDF

$$\text{CDF}(R|\vec{\nu}^{\text{ center}}, \vec{\nu}^{\text{ surround}}, w_{\text{MI}}) = \begin{cases} \Phi(R; b + \vec{\nu}^{\text{ center}} - w_{\text{MI}}\vec{\nu}^{\text{ surround}}, \\ \sigma^2_{\text{motor}} + \sigma^2_{\text{center}} + w^2_{\text{MI}}\sigma^2_{\text{surround}}) & w_{\text{MI}} \geq 0 \\ \Phi(R; b + (1 + w_{\text{MI}})\vec{\nu}^{\text{ center}} - w_{\text{MI}}\vec{\nu}^{\text{ surround}}, \\ \sigma^2_{\text{motor}} + (1 + w_{\text{MI}})^2\sigma^2_{\text{center}} + w^2_{\text{MI}}\sigma^2_{\text{surround}}) & w_{\text{MI}} \leq 0 \end{cases} \tag{S2}$$

where $\Phi$ denotes the normal CDF. The variance over reports in the stationary surround condition, given by $\text{Var}(R_{\text{stationary}}) = \sigma^2_{\text{motor}} + \sigma^2_{\text{center}}$ is an upper bound on the $\sigma^2_{\text{center}}$ (as $\sigma^2_{\text{motor}} \geq 0$). This allowed us to parameterize $\sigma^2_{\text{center}} = \text{Var}(R_{\text{stationary}})c_{\text{center}}$ and $\sigma^2_{\text{surround}} = \text{Var}(R_{\text{stationary}})c_{\text{surround}}$ where $0 \leq c_{\text{center}}, c_{\text{surround}} \leq 1$. In order to estimate $c_{\text{center}}$, $c_{\text{surround}}$, and $\pi^i_{\text{MI}}$, we minimized the L2 norm between the predicted CDF over the reported directions and the empirically estimated CDF in the moving surround condition. The predicted CDF can be expanded as a mixture of the CDF predicted for each $w_{\text{MI}}$ weighted by the corresponding probability:

$$\text{CDF}(R|\vec{\nu}^{\text{ center}}, \vec{\nu}^{\text{ surround}}) = \sum_{i=1}^{33} \text{CDF}(R|\vec{\nu}^{\text{ center}}, \vec{\nu}^{\text{ surround}}, w^i_{\text{MI}})\pi^i_{\text{MI}}\Delta w_{\text{MI}}. \tag{S3}$$

$w^i_{\text{MI}}$ are the bin centers separated by $\Delta w_{\text{MI}}$. We assumed a weakly informative lognormal prior over $\sigma^2_{\text{center}}$ and $\sigma^2_{\text{surround}}$ (with logarithmic mean $-3.75$ and standard deviation 1.15). We also added two costs on $\pi^i_{\text{MI}}$ as regularization to ensure convergence: (a) L1 regularization penalizing the sum of absolute values of $\pi^i_{\text{MI}}$ that ensured that $\pi^i_{\text{MI}}$ corresponding to non-required regions of $w^i_{\text{MI}}$ went to zero, and (b) regularization on the curvature (second derivative) to ensure smooth distributions over $w^i_{\text{MI}}$. The regularization constants $\lambda_1$ and $\lambda_2$ corresponding to the two terms of the regularization were estimated by cross validation on synthetic ground truth data.

#### B Possible structures for $n$ moving elements

The causal structures for $n$ moving elements can be decomposed into $T(n)$ grouping trees. Each grouping tree indicates which elements are grouped together. If $C_i$ denotes the number of nodes in the grouping tree $i$, then the number of structures corresponding to a grouping tree is $2^{C_i}$ since each node of the grouping tree contains a relative velocity that is either zero or non-zero, contributing 2 structures.

To enumerate the number of grouping trees, we construct an $n \times n$ grouping tree matrix $\mathcal{T}$ in which $\mathcal{T}_{i,j}$ denotes the number of grouping trees of depth $i$ containing $j$ nodes at top level. The sum of all elements in $\mathcal{T}$ gives the total number of grouping trees. If all elements are inferred to be moving independently, then the number of nodes at level 1 are $n$, i.e. $\mathcal{T}_{1,n} = 1$. The values of the matrix at subsequent levels can be derived recursively using the relation

$$\mathcal{T}_{i,j} = \sum_{k=j+1}^{n-i+1} \mathcal{T}_{i-1,k} \begin{Bmatrix} k \\ j \end{Bmatrix} \tag{S4}$$

wherein $\left\{{n \atop k}\right\}$ are Stirling numbers of the second kind. The recursion counts the number of ways to partition $k$ nodes in the previous level into $j$ nodes at the current level. The number of grouping trees, $T(n)$, for $n = 1..5$ moving elements are $1, 2, 8, 64, 872$, respectively.

To derive the number of nodes in each grouping tree, we compute the node matrix $\mathcal{C}$ in which $\mathcal{C}_{i,j}$ denotes the number of unique nodes corresponding to the grouping trees in $\mathcal{T}_{i,j}$. Given that $\mathcal{C}_{1,n} = n$, the other elements can be computed recursively as

$$\mathcal{C}_{i,j} = \sum_{k=j+1}^{n-i+1} \mathcal{C}_{i-1,k} + j \tag{S5}$$

This however overestimates the number of nodes as non-grouped elements at lower levels are assigned into a new singleton group at each higher level. The number of $a$ singleton branches of depth 1 (denoted as $\mathcal{L}_{i,j,a,1}$) can be estimated as

$$\mathcal{L}_{i,j,a,1} = \sum_{k=j+1}^{n-i+2} \mathcal{T}_{i-1,k} \binom{k}{a} S_2(k-a, j-a) \tag{S6}$$

in which $S_r(n,k)$ denote the r-associated Stirling numbers of the second kind. The resulting number of structures for $n = 1..5$ moving elements are $2, 12, 264, 25392, 8281376$, respectively.

While we have derived expressions for counting the number of structures, listing them and deriving inference equations for each structure individually is intractable. For small $n$, we can instead consider the largest grouping tree which has $C = \frac{N(N+1)}{2}$ nodes and derive inference equation for this single tree but gating each node based on the structures in occurs in. This partitioning of the nodes independent of the grouping trees requires deriving a single inference model which is what we use for our model fitting for 2 moving elements.

### C Inference in the hierarchical causal inference model

In this section, we derive the expressions for posteriors over the inferred latents in our causal inference model. Eq. 9 in the main text can be expanded in the context of our model as

$$\begin{aligned} p(\vec{v}^{\text{center}}_{\text{percept}} | \vec{o}) = &\sum_{c_1=\{0,1\}} \sum_{c_2=\{0,1\}} \sum_{c_3=\{0,1\}} \sum_{c_4=\{0,1\}} \\ &p(C^{\text{center}} = c_1, C^{\text{surround}} = c_2, C^{\text{group}} = c_3, S^{\text{center,surround}} = c_4 | \vec{o}) \\ &p(\vec{v}^{\text{center}}_{\text{percept}} | \vec{o}, C^{\text{center}} = c_1, C^{\text{surround}} = c_2, C^{\text{group}} = c_3, S^{\text{center,surround}} = c_4). \end{aligned} \tag{S7}$$

where $w_i(\vec{o}_j) = p(C^{\text{center}} = c_1, C^{\text{surround}} = c_2, C^{\text{group}} = c_3, S^{\text{center,surround}} = c_4 | \vec{o})$, $\mu_i(\vec{o}_j)$ and $\sigma_i^2$ are the mean and variance of $p(\vec{v}^{\text{center}}_{\text{percept}} | \vec{o}, C^{\text{center}} = c_1, C^{\text{surround}} = c_2, C^{\text{group}} = c_3, S^{\text{center,surround}} = c_4)$ respectively.

To get the required expression for Eq. S7, it is sufficient to derive expressions for

$$p(C^{\text{ center}}, S^{\text{center,surround}}, C^{\text{group}}, C^{\text{ surround}} | \vec{o}^{\text{ center}}, \vec{o}^{\text{ surround}}) \tag{S8}$$

$$p(\vec{v}^{\text{group}}_{\text{relative}} | \vec{o}^{\text{ center}}, \vec{o}^{\text{ surround}}, C^{\text{ center}}, S^{\text{center,surround}}, C^{\text{group}}, C^{\text{ surround}}) \tag{S9}$$

$$p(\vec{v}^{\text{center}}_{\text{relative}} | \vec{o}^{\text{ center}}, \vec{o}^{\text{ surround}}, C^{\text{ center}}, S^{\text{center,surround}}, C^{\text{group}}, C^{\text{ surround}}) \tag{S10}$$

as

$$\begin{aligned} &p(\vec{v}^{\text{center}}_{\text{percept}} | \vec{o}^{\text{ center}}, \vec{o}^{\text{ surround}}, C^{\text{ center}} = 0, S^{\text{center,surround}} = 1, C^{\text{group}} = 1, C^{\text{ surround}} \in \{0,1\}) = \\ &\quad p(\vec{v}^{\text{group}}_{\text{relative}} | \vec{o}^{\text{ center}}, \vec{o}^{\text{ surround}}, C^{\text{ center}} = 0, S^{\text{center,surround}} = 1, C^{\text{group}} = 1, C^{\text{ surround}} \in \{0,1\}) \end{aligned} \tag{S11}$$

$$
\begin{aligned}
&p(\vec{v}^{\text{center}}_{\text{percept}}|\vec{o}^{\text{ center}}, \vec{o}^{\text{ surround}}, C^{\text{ center}} = 1, S^{\text{center,surround}} \in \{0,1\}, C^{\text{group}} \in \{0,1\}, C^{\text{ surround}} \in \{0,1\}) = \\
&\quad p(\vec{v}^{\text{center}}_{\text{relative}}|\vec{o}^{\text{ center}}, \vec{o}^{\text{ surround}}, C^{\text{ center}} = 1, S^{\text{center,surround}} \in \{0,1\}, C^{\text{group}} \in \{0,1\}, C^{\text{ surround}} \in \{0,1\})
\end{aligned}
\tag{S12}
$$

The joint distribution over the latents and observations for a given structure can be expanded in terms of their definitions (main text)

$$
\begin{aligned}
&p(\vec{o}^{\text{ center}}, \vec{o}^{\text{ surround}}, \vec{v}^{\text{group}}_{\text{relative}}, \vec{v}^{\text{center}}_{\text{relative}}, \vec{v}^{\text{surround}}_{\text{relative}}, \vec{v}^{\text{group}}|C^{\text{ center}}, S^{\text{center,surround}}, C^{\text{group}}, C^{\text{ surround}}) = \\
&\quad \mathcal{N}(\vec{o}^{\text{ center}} - \vec{v}^{\text{center}}_{\text{relative}}; S^{\text{center,surround}}\vec{v}^{\text{group}}, \sigma^2_{\text{center}} + \sigma^2_{\Delta}) \\
&\quad \mathcal{N}(\vec{o}^{\text{ surround}} - \vec{v}^{\text{surround}}_{\text{relative}}; S^{\text{center,surround}}\vec{v}^{\text{group}}, \sigma^2_{\text{surround}} + \sigma^2_{\Delta}) \\
&\quad \mathcal{N}(\vec{v}^{\text{group}}; \vec{v}^{\text{group}}_{\text{relative}}, \sigma^2_{\Delta})\mathcal{N}(\vec{v}^{\text{group}}_{\text{relative}}; 0, C^{\text{group}}\sigma^2_{\text{prior}}) \\
&\quad \mathcal{N}(\vec{v}^{\text{center}}_{\text{relative}}; 0, C^{\text{ center}}\sigma^2_{\text{prior}})\mathcal{N}(\vec{v}^{\text{surround}}_{\text{relative}}; 0, C^{\text{ surround}}\sigma^2_{\text{prior}})
\end{aligned}
\tag{S13}
$$

We have assumed the inferred $\vec{v}^{\text{ world}} = 0$ since our stimuli in the experiment are local moving patches and are unlikely to introduce non-zero self motion velocities. Marginalizing out the center and surround velocities

$$
\begin{aligned}
&p(\vec{o}^{\text{ center}}, \vec{o}^{\text{ surround}}, \vec{v}^{\text{group}}_{\text{relative}}, \vec{v}^{\text{group}}|C^{\text{ center}}, S^{\text{center,surround}}, C^{\text{group}}, C^{\text{ surround}}) = \\
&\quad \mathcal{N}(\vec{o}^{\text{ center}}; S^{\text{center,surround}}\vec{v}^{\text{group}}, \sigma^2_{\text{center}} + \sigma^2_{\Delta} + C^{\text{ center}}\sigma^2_{\text{prior}}) \\
&\quad \mathcal{N}(\vec{o}^{\text{ surround}}; S^{\text{center,surround}}\vec{v}^{\text{group}}, \sigma^2_{\text{surround}} + \sigma^2_{\Delta} + C^{\text{ surround}}\sigma^2_{\text{prior}}) \\
&\quad \mathcal{N}(\vec{v}^{\text{group}}; \vec{v}^{\text{group}}_{\text{relative}}, \sigma^2_{\Delta})\mathcal{N}(\vec{v}^{\text{group}}_{\text{relative}}; 0, C^{\text{group}}\sigma^2_{\text{prior}})
\end{aligned}
\tag{S14}
$$

By defining

$$
\sigma_5^2 = \sigma^2_{\text{center}} + \sigma^2_{\text{surround}} + 2\sigma^2_{\Delta} + C^{\text{ center}}\sigma^2_{\text{prior}} + C^{\text{ surround}}\sigma^2_{\text{prior}}
\tag{S15}
$$

$$
\gamma_5 = \frac{\sigma^2_{\text{surround}} + \sigma^2_{\Delta} + C^{\text{ surround}}\sigma^2_{\text{prior}}}{\sigma_5^2}
\tag{S16}
$$

we can apply the formula for product of Gaussian pdf to get

$$
\begin{aligned}
&p(\vec{o}^{\text{ center}}, \vec{o}^{\text{ surround}}, \vec{v}^{\text{group}}_{\text{relative}}|C^{\text{ center}}, S^{\text{center,surround}}, C^{\text{group}}, C^{\text{ surround}}) = \mathcal{N}(\vec{o}^{\text{ center}}; \vec{o}^{\text{ surround}}, \sigma_5^2) \\
&\quad \mathcal{N}(\vec{o}^{\text{ center}}\gamma_5 + \vec{o}^{\text{ surround}}(1-\gamma_5); S^{\text{center,surround}}\vec{v}^{\text{group}}_{\text{relative}}, \sigma_5^2\gamma_5(1-\gamma_5) + S^{\text{center,surround}}\sigma^2_{\Delta}) \\
&\quad \mathcal{N}(\vec{v}^{\text{group}}_{\text{relative}}; 0, C^{\text{group}}\sigma^2_{\text{prior}})
\end{aligned}
\tag{S17}
$$

Similarly by defining

$$
\sigma_6^2 = \sigma_5^2\gamma_5(1-\gamma_5) + S^{\text{center,surround}}(\sigma^2_{\Delta} + C^{\text{group}}\sigma^2_{\text{prior}})
\tag{S18}
$$

$$
\gamma_6 = \frac{S^{\text{center,surround}}C^{\text{group}}\sigma^2_{\text{prior}}}{\sigma_6^2}
\tag{S19}
$$

we can apply the product of Gaussian pdf to get

$$p(\vec{o}^{\,\text{center}},\vec{o}^{\,\text{surround}},\vec{v}^{\text{group}}_{\text{relative}}|C^{\,\text{center}},S^{\text{center,surround}},C^{\text{group}},C^{\,\text{surround}}) = \\ \mathcal{N}(\vec{o}^{\,\text{center}};\vec{o}^{\,\text{surround}},\sigma_5^2)\mathcal{N}(\vec{o}^{\,\text{center}}\gamma_5+\vec{o}^{\,\text{surround}}(1-\gamma_5);0,\sigma_6^2) \\ \mathcal{N}(\vec{v}^{\text{group}}_{\text{relative}};\vec{o}^{\,\text{center}}\gamma_5\gamma_6+\vec{o}^{\,\text{surround}}(1-\gamma_5)\gamma_6,C^{\text{group}}\sigma^2_{\text{prior}}(1-S^{\text{center,surround}}\gamma_6)) \tag{S20}$$

We can substitute the above equation into the likelihood for the posterior over the group relative velocity. By marginalizing the group relative velocity, we can also substitute the above equation to the posterior over the different causal structures

$$p(\vec{v}^{\text{group}}_{\text{relative}}|\vec{o}^{\,\text{center}},\vec{o}^{\,\text{surround}},C^{\,\text{center}},S^{\text{center,surround}},C^{\text{group}},C^{\,\text{surround}}) = \\ \mathcal{N}(\vec{v}^{\text{group}}_{\text{relative}};\vec{o}^{\,\text{center}}\gamma_5\gamma_6+\vec{o}^{\,\text{surround}}(1-\gamma_5)\gamma_6,C^{\text{group}}\sigma^2_{\text{prior}}(1-S^{\text{center,surround}}\gamma_6)) \tag{S21}$$

$$p(C^{\,\text{center}},S^{\text{center,surround}},C^{\text{group}},C^{\,\text{surround}}|\vec{o}^{\,\text{center}},\vec{o}^{\,\text{surround}}) = \\ \mathcal{N}(\vec{o}^{\,\text{center}};\vec{o}^{\,\text{surround}},\sigma_5^2)\mathcal{N}(\vec{o}^{\,\text{center}}\gamma_5+\vec{o}^{\,\text{surround}}(1-\gamma_5);0,\sigma_6^2) \\ (1-\alpha)^{(C^{\,\text{center}}+C^{\text{group}}+C^{\,\text{surround}})}(\alpha)^{(3-C^{\,\text{center}}-C^{\text{group}}-C^{\,\text{surround}})}\beta_{\text{gr}}^{S^{\text{center,surround}}}(1-\beta_{\text{gr}})^{(1-S^{\text{center,surround}})} \tag{S22}$$

To get the posterior over the center relative velocity, we start with joint distributions over the latents for each causal structures and repeatedly apply the Gaussian pdf products and marginalize out the variables other than $\vec{v}^{\text{center}}_{\text{relative}}$

$$p(\vec{o}^{\,\text{center}},\vec{o}^{\,\text{surround}},\vec{v}^{\text{group}}_{\text{relative}},\vec{v}^{\text{center}}_{\text{relative}},\vec{v}^{\text{surround}}_{\text{relative}},\vec{v}^{\text{group}}|C^{\,\text{center}},S^{\text{center,surround}},C^{\text{group}},C^{\,\text{surround}}) = \\ \mathcal{N}(\vec{o}^{\,\text{center}}-\vec{v}^{\text{center}}_{\text{relative}};S^{\text{center,surround}}\vec{v}^{\text{group}},\sigma^2_{\text{center}}+\sigma^2_{\Delta}) \\ \mathcal{N}(\vec{o}^{\,\text{surround}}-\vec{v}^{\text{surround}}_{\text{relative}};S^{\text{center,surround}}\vec{v}^{\text{group}},\sigma^2_{\text{surround}}+\sigma^2_{\Delta}) \\ \mathcal{N}(\vec{v}^{\text{group}};\vec{v}^{\text{group}}_{\text{relative}},\sigma^2_{\Delta})\mathcal{N}(\vec{v}^{\text{group}}_{\text{relative}};0,C^{\text{group}}\sigma^2_{\text{prior}})\mathcal{N}(\vec{v}^{\text{center}}_{\text{relative}};0,C^{\,\text{center}}\sigma^2_{\text{prior}})\mathcal{N}(\vec{v}^{\text{surround}}_{\text{relative}};0,C^{\,\text{surround}}\sigma^2_{\text{prior}}) \tag{S23}$$

By defining

$$\sigma_1^2=\sigma^2_{\text{surround}}+\sigma^2_{\text{center}}+2\sigma^2_{\Delta} \tag{S24}$$

$$\gamma_1=\frac{\sigma^2_{\text{surround}}+\sigma^2_{\Delta}}{\sigma_1^2} \tag{S25}$$

we can apply the formula for product of Gaussian pdf to get

$$p(\vec{o}^{\,\text{center}},\vec{o}^{\,\text{surround}},\vec{v}^{\text{group}}_{\text{relative}},\vec{v}^{\text{center}}_{\text{relative}},\vec{v}^{\text{surround}}_{\text{relative}}|C^{\,\text{center}},S^{\text{center,surround}},C^{\text{group}},C^{\,\text{surround}}) = \\ \mathcal{N}(\vec{o}^{\,\text{center}}-\vec{v}^{\text{center}}_{\text{relative}};\vec{o}^{\,\text{surround}}-\vec{v}^{\text{surround}}_{\text{relative}},\sigma_1^2) \\ \mathcal{N}(S^{\text{center,surround}}\vec{v}^{\text{group}}_{\text{relative}};(\vec{o}^{\,\text{center}}-\vec{v}^{\text{center}}_{\text{relative}})\gamma_1+(\vec{o}^{\,\text{surround}}-\vec{v}^{\text{surround}}_{\text{relative}})(1-\gamma_1),\sigma_1^2\gamma_1(1-\gamma_1)+S^{\text{center,surround}}\sigma^2_{\Delta}) \\ \mathcal{N}(\vec{v}^{\text{group}}_{\text{relative}};0,C^{\text{group}}\sigma^2_{\text{prior}})\mathcal{N}(\vec{v}^{\text{center}}_{\text{relative}};0,C^{\,\text{center}}\sigma^2_{\text{prior}})\mathcal{N}(\vec{v}^{\text{surround}}_{\text{relative}};0,C^{\,\text{surround}}\sigma^2_{\text{prior}}) \tag{S26}$$

By marginalizing the group relative velocity

$$\begin{gathered} p(\vec{o}^{\,\text{center}}, \vec{o}^{\,\text{surround}}, \vec{v}^{\text{center}}_{\text{relative}}, \vec{v}^{\text{surround}}_{\text{relative}} | C^{\,\text{center}}, S^{\text{center,surround}}, C^{\text{group}}, C^{\,\text{surround}}) = \\ \mathcal{N}(\vec{o}^{\,\text{center}} - \vec{v}^{\text{center}}_{\text{relative}}; \vec{o}^{\,\text{surround}} - \vec{v}^{\text{surround}}_{\text{relative}}, \sigma_1^2) \\ \mathcal{N}(0; (\vec{o}^{\,\text{center}} - \vec{v}^{\text{center}}_{\text{relative}})\gamma_1 + (\vec{o}^{\,\text{surround}} - \vec{v}^{\text{surround}}_{\text{relative}})(1-\gamma_1), \sigma_1^2\gamma_1(1-\gamma_1) + S^{\text{center,surround}}(\sigma_\Delta^2 + C^{\text{group}}\sigma^2_{\text{prior}})) \\ \mathcal{N}(\vec{v}^{\text{center}}_{\text{relative}}; 0, C^{\,\text{center}}\sigma^2_{\text{prior}})\mathcal{N}(\vec{v}^{\text{surround}}_{\text{relative}}; 0, C^{\,\text{surround}}\sigma^2_{\text{prior}}) \end{gathered} \tag{S27}$$

Rearraging the different terms

$$\begin{gathered} p(\vec{o}^{\,\text{center}}, \vec{o}^{\,\text{surround}}, \vec{v}^{\text{center}}_{\text{relative}}, \vec{v}^{\text{surround}}_{\text{relative}} | C^{\,\text{center}}, S^{\text{center,surround}}, C^{\text{group}}, C^{\,\text{surround}}) = \\ \mathcal{N}(\vec{v}^{\text{surround}}_{\text{relative}}; \vec{o}^{\,\text{surround}} - \vec{o}^{\,\text{center}} + \vec{v}^{\text{center}}_{\text{relative}}, \sigma_1^2) \\ \mathcal{N}((1-\gamma_1)\vec{v}^{\text{surround}}_{\text{relative}}; (\vec{o}^{\,\text{center}} - \vec{v}^{\text{center}}_{\text{relative}})\gamma_1 + \vec{o}^{\,\text{surround}}(1-\gamma_1), \sigma_1^2\gamma_1(1-\gamma_1) + S^{\text{center,surround}}(\sigma_\Delta^2 + C^{\text{group}}\sigma^2_{\text{prior}})) \\ \mathcal{N}(\vec{v}^{\text{center}}_{\text{relative}}; 0, C^{\,\text{center}}\sigma^2_{\text{prior}})\mathcal{N}(\vec{v}^{\text{surround}}_{\text{relative}}; 0, C^{\,\text{surround}}\sigma^2_{\text{prior}}) \end{gathered} \tag{S28}$$

$$\begin{gathered} p(\vec{o}^{\,\text{center}}, \vec{o}^{\,\text{surround}}, \vec{v}^{\text{center}}_{\text{relative}}, \vec{v}^{\text{surround}}_{\text{relative}} | C^{\,\text{center}}, S^{\text{center,surround}}, C^{\text{group}}, C^{\,\text{surround}}) = \\ \mathcal{N}(\vec{v}^{\text{surround}}_{\text{relative}}; \vec{o}^{\,\text{surround}} - \vec{o}^{\,\text{center}} + \vec{v}^{\text{center}}_{\text{relative}}, \sigma_1^2)\frac{1}{(1-\gamma_1)^2} \\ \mathcal{N}(\vec{v}^{\text{surround}}_{\text{relative}}; (\vec{o}^{\,\text{center}} - \vec{v}^{\text{center}}_{\text{relative}})\frac{\gamma_1}{(1-\gamma_1)} + \vec{o}^{\,\text{surround}}, \frac{\sigma_1^2\gamma_1(1-\gamma_1) + S^{\text{center,surround}}(\sigma_\Delta^2 + C^{\text{group}}\sigma^2_{\text{prior}})}{(1-\gamma_1)^2}) \\ \mathcal{N}(\vec{v}^{\text{center}}_{\text{relative}}; 0, C^{\,\text{center}}\sigma^2_{\text{prior}})\mathcal{N}(\vec{v}^{\text{surround}}_{\text{relative}}; 0, C^{\,\text{surround}}\sigma^2_{\text{prior}}) \end{gathered} \tag{S29}$$

By defining

$$\sigma_2^2 = \sigma_1^2 + \frac{\sigma_1^2\gamma_1(1-\gamma_1) + S^{\text{center,surround}}(\sigma_\Delta^2 + C^{\text{group}}\sigma^2_{\text{prior}})}{(1-\gamma_1)^2} \tag{S30}$$

$$\gamma_2 = 1 - \frac{\sigma_1^2}{\sigma_2^2} \tag{S31}$$

we can apply the formula for product of Gaussian pdf and marginalizing the surround relative velocity to get

$$\begin{gathered} p(\vec{o}^{\,\text{center}}, \vec{o}^{\,\text{surround}}, \vec{v}^{\text{center}}_{\text{relative}} | C^{\,\text{center}}, S^{\text{center,surround}}, C^{\text{group}}, C^{\,\text{surround}}) = \\ \frac{1}{(1-\gamma_1)^2}\mathcal{N}((\vec{o}^{\,\text{center}} - \vec{v}^{\text{center}}_{\text{relative}})\frac{\gamma_1}{(1-\gamma_1)} + \vec{o}^{\,\text{surround}}; \vec{o}^{\,\text{surround}} - \vec{o}^{\,\text{center}} + \vec{v}^{\text{center}}_{\text{relative}}, \sigma_2^2) \\ \mathcal{N}(0; (\vec{o}^{\,\text{surround}} - \vec{o}^{\,\text{center}} + \vec{v}^{\text{center}}_{\text{relative}})\gamma_2 + ((\vec{o}^{\,\text{center}} - \vec{v}^{\text{center}}_{\text{relative}})\frac{\gamma_1}{(1-\gamma_1)} + \vec{o}^{\,\text{surround}})(1-\gamma_2), \sigma_2^2\gamma_2(1-\gamma_2) + C^{\,\text{surround}}\sigma^2_{\text{prior}}) \\ \mathcal{N}(\vec{v}^{\text{center}}_{\text{relative}}; 0, C^{\,\text{center}}\sigma^2_{\text{prior}}) \end{gathered} \tag{S32}$$

By rearranging terms

$$p(\vec{o}^{\,\text{center}}, \vec{o}^{\,\text{surround}}, \vec{v}^{\text{center}}_{\text{relative}} | C^{\,\text{center}}, S^{\text{center,surround}}, C^{\text{group}}, C^{\,\text{surround}}) =$$
$$(1-\gamma_1)^2 \mathcal{N}(\vec{o}^{\,\text{center}}\gamma_1 - \vec{v}^{\text{center}}_{\text{relative}}\gamma_1 + \vec{o}^{\,\text{surround}}(1-\gamma_1); \vec{o}^{\,\text{surround}}(1-\gamma_1) - \vec{o}^{\,\text{center}}(1-\gamma_1) + \vec{v}^{\text{center}}_{\text{relative}}(1-\gamma_1), \sigma_2^2(1-\gamma_1)^2)$$
$$\mathcal{N}(0; (\vec{o}^{\,\text{surround}} - \vec{o}^{\,\text{center}} + \vec{v}^{\text{center}}_{\text{relative}})\gamma_2(1-\gamma_1) + (\vec{o}^{\,\text{center}}\gamma_1 - \vec{v}^{\text{center}}_{\text{relative}}\gamma_1 + \vec{o}^{\,\text{surround}}(1-\gamma_1))(1-\gamma_2), \ldots$$
$$\sigma_1^2\gamma_2(1-\gamma_1)^2 + C^{\,\text{surround}}\sigma^2_{\text{prior}}(1-\gamma_1)^2)$$
$$\mathcal{N}(\vec{v}^{\text{center}}_{\text{relative}}; 0, C^{\,\text{center}}\sigma^2_{\text{prior}}) \tag{S33}$$

$$p(\vec{o}^{\,\text{center}}, \vec{o}^{\,\text{surround}}, \vec{v}^{\text{center}}_{\text{relative}} | C^{\,\text{center}}, S^{\text{center,surround}}, C^{\text{group}}, C^{\,\text{surround}}) =$$
$$(1-\gamma_1)^2 \mathcal{N}(\vec{v}^{\text{center}}_{\text{relative}}; \vec{o}^{\,\text{center}}, \sigma_2^2(1-\gamma_1)^2)$$
$$\mathcal{N}((\gamma_1-\gamma_2)\vec{v}^{\text{center}}_{\text{relative}}; (\gamma_1-\gamma_2)\vec{o}^{\,\text{center}} + \vec{o}^{\,\text{surround}}(1-\gamma_1), \sigma_1^2\gamma_2(1-\gamma_1)^2 + C^{\,\text{surround}}\sigma^2_{\text{prior}}(1-\gamma_1)^2)$$
$$\mathcal{N}(\vec{v}^{\text{center}}_{\text{relative}}; 0, C^{\,\text{center}}\sigma^2_{\text{prior}}) \tag{S34}$$

$$p(\vec{o}^{\,\text{center}}, \vec{o}^{\,\text{surround}}, \vec{v}^{\text{center}}_{\text{relative}} | C^{\,\text{center}}, S^{\text{center,surround}}, C^{\text{group}}, C^{\,\text{surround}}) =$$
$$(1-\gamma_1)^2 \mathcal{N}((\gamma_1-\gamma_2)\vec{v}^{\text{center}}_{\text{relative}}; (\gamma_1-\gamma_2)\vec{o}^{\,\text{center}} + \vec{o}^{\,\text{surround}}(1-\gamma_1), \sigma_1^2\gamma_2(1-\gamma_1)^2 + C^{\,\text{surround}}\sigma^2_{\text{prior}}(1-\gamma_1)^2)$$
$$\mathcal{N}(\vec{v}^{\text{center}}_{\text{relative}}; \vec{o}^{\,\text{center}}, \sigma_2^2(1-\gamma_1)^2)\mathcal{N}(\vec{v}^{\text{center}}_{\text{relative}}; 0, C^{\,\text{center}}\sigma^2_{\text{prior}}) \tag{S35}$$

$$p(\vec{o}^{\,\text{center}}, \vec{o}^{\,\text{surround}}, \vec{v}^{\text{center}}_{\text{relative}} | C^{\,\text{center}}, S^{\text{center,surround}}, C^{\text{group}}, C^{\,\text{surround}}) = \frac{(1-\gamma_1)^2}{(\gamma_1-\gamma_2)^2}$$
$$\mathcal{N}(\vec{v}^{\text{center}}_{\text{relative}}; \vec{o}^{\,\text{center}} + \vec{o}^{\,\text{surround}}\frac{(1-\gamma_1)}{(\gamma_1-\gamma_2)}, \sigma_1^2\gamma_2\frac{(1-\gamma_1)^2}{(\gamma_1-\gamma_2)^2} + C^{\,\text{surround}}\sigma^2_{\text{prior}}\frac{(1-\gamma_1)^2}{(\gamma_1-\gamma_2)^2})$$
$$\mathcal{N}(\vec{v}^{\text{center}}_{\text{relative}}; \vec{o}^{\,\text{center}}, \sigma_2^2(1-\gamma_1)^2)\mathcal{N}(\vec{v}^{\text{center}}_{\text{relative}}; 0, C^{\,\text{center}}\sigma^2_{\text{prior}}) \tag{S36}$$

By defining

$$\sigma_3^2 = (\sigma_1^2\gamma_2 + C^{\,\text{surround}}\sigma^2_{\text{prior}} + \sigma_2^2(\gamma_1-\gamma_2)^2)(1-\gamma_1)^2 \tag{S37}$$

$$\gamma_3 = \frac{\sigma_2^2(1-\gamma_1)^2(\gamma_1-\gamma_2)^2}{\sigma_3^2} \tag{S38}$$

we can apply the formula for product of Gaussian pdf

$$p(\vec{o}^{\,\text{center}}, \vec{o}^{\,\text{surround}}, \vec{v}^{\text{center}}_{\text{relative}} | C^{\,\text{center}}, S^{\text{center,surround}}, C^{\text{group}}, C^{\,\text{surround}}) =$$
$$\frac{(1-\gamma_1)^2}{(\gamma_1-\gamma_2)^2}\mathcal{N}(\vec{o}^{\,\text{center}}; \vec{o}^{\,\text{center}} + \vec{o}^{\,\text{surround}}\frac{(1-\gamma_1)}{(\gamma_1-\gamma_2)}, \frac{\sigma_3^2}{(\gamma_1-\gamma_2)^2})$$
$$\mathcal{N}(\vec{v}^{\text{center}}_{\text{relative}}; \vec{o}^{\,\text{center}} + \vec{o}^{\,\text{surround}}\frac{(1-\gamma_1)}{(\gamma_1-\gamma_2)}S^{\text{center,surround}}\gamma_3, \sigma_2^2(1-\gamma_1)^2(1-S^{\text{center,surround}}\gamma_3))$$
$$\mathcal{N}(\vec{v}^{\text{center}}_{\text{relative}}; 0, C^{\,\text{center}}\sigma^2_{\text{prior}}) \tag{S39}$$

By rearranging terms

$$
\begin{aligned}
p(\vec{o}^{\,\text{center}}, \vec{o}^{\,\text{surround}}, \vec{v}^{\text{center}}_{\text{relative}} | C^{\,\text{center}}, S^{\text{center,surround}}, C^{\text{group}}, C^{\,\text{surround}}) = \\
(1-\gamma_1)^2 \mathcal{N}((\gamma_1-\gamma_2)\vec{o}^{\,\text{center}} + \vec{o}^{\,\text{surround}}(1-\gamma_1); (\gamma_1-\gamma_2)\vec{o}^{\,\text{center}}, \sigma_3^2) \\
\mathcal{N}(\vec{v}^{\text{center}}_{\text{relative}}; \vec{o}^{\,\text{center}} + \vec{o}^{\,\text{surround}} \frac{(1-\gamma_1)}{(\gamma_1-\gamma_2)} S^{\text{center,surround}}\gamma_3, \sigma_2^2(1-\gamma_1)^2(1-S^{\text{center,surround}}\gamma_3)) \\
\mathcal{N}(\vec{v}^{\text{center}}_{\text{relative}}; 0, C^{\,\text{center}}\sigma^2_{\text{prior}})
\end{aligned}
\tag{S40}
$$

We can expand and simplify some of the terms in the pdf to define a gain $g$

$$\sigma_2^2 = \frac{\sigma_1^2}{(1-\gamma_1)} + \frac{S^{\text{center,surround}}(\sigma_\Delta^2 + C^{\text{group}}\sigma^2_{\text{prior}})}{(1-\gamma_1)^2} \tag{S41}$$

$$\sigma_2^2 = \frac{\sigma_1^2(1-\gamma_1) + S^{\text{center,surround}}(\sigma_\Delta^2 + C^{\text{group}}\sigma^2_{\text{prior}})}{(1-\gamma_1)^2} \tag{S42}$$

$$\sigma_2^2 - \sigma_1^2 = \frac{\sigma_1^2(1-\gamma_1)\gamma_1 + S^{\text{center,surround}}(\sigma_\Delta^2 + C^{\text{group}}\sigma^2_{\text{prior}})}{(1-\gamma_1)^2} \tag{S43}$$

$$\gamma_2 = \frac{\sigma_1^2\gamma_1(1-\gamma_1) + S^{\text{center,surround}}(\sigma_\Delta^2 + C^{\text{group}}\sigma^2_{\text{prior}})}{\sigma_1^2(1-\gamma_1) + S^{\text{center,surround}}(\sigma_\Delta^2 + C^{\text{group}}\sigma^2_{\text{prior}})} \tag{S44}$$

$$\gamma_1 - \gamma_2 = \gamma_1 - \frac{\sigma_1^2\gamma_1(1-\gamma_1) + S^{\text{center,surround}}(\sigma_\Delta^2 + C^{\text{group}}\sigma^2_{\text{prior}})}{\sigma_1^2(1-\gamma_1) + S^{\text{center,surround}}(\sigma_\Delta^2 + C^{\text{group}}\sigma^2_{\text{prior}})} \tag{S45}$$

$$\gamma_1 - \gamma_2 = \frac{S^{\text{center,surround}}\gamma_1(\sigma_\Delta^2 + C^{\text{group}}\sigma^2_{\text{prior}}) - S^{\text{center,surround}}(\sigma_\Delta^2 + C^{\text{group}}\sigma^2_{\text{prior}})}{\sigma_1^2(1-\gamma_1) + S^{\text{center,surround}}(\sigma_\Delta^2 + C^{\text{group}}\sigma^2_{\text{prior}})} \tag{S46}$$

$$\gamma_1 - \gamma_2 = -\frac{S^{\text{center,surround}}(\sigma_\Delta^2 + C^{\text{group}}\sigma^2_{\text{prior}})(1-\gamma_1)}{\sigma_1^2(1-\gamma_1) + S^{\text{center,surround}}(\sigma_\Delta^2 + C^{\text{group}}\sigma^2_{\text{prior}})} \tag{S47}$$

$$(\gamma_1 - \gamma_2)^2 = \frac{S^{\text{center,surround}}(\sigma_\Delta^2 + C^{\text{group}}\sigma^2_{\text{prior}})^2(1-\gamma_1)^2}{[\sigma_1^2(1-\gamma_1) + S^{\text{center,surround}}(\sigma_\Delta^2 + C^{\text{group}}\sigma^2_{\text{prior}})]^2} \tag{S48}$$

$$\sigma_2^2(\gamma_1 - \gamma_2)^2 = \frac{S^{\text{center,surround}}(\sigma_\Delta^2 + C^{\text{group}}\sigma^2_{\text{prior}})^2}{[\sigma_1^2(1-\gamma_1) + S^{\text{center,surround}}(\sigma_\Delta^2 + C^{\text{group}}\sigma^2_{\text{prior}})]} \tag{S49}$$

$$\frac{1}{(\sigma_1^2\gamma_2 + C^{\,\text{surround}}\sigma^2_{\text{prior}} + \sigma_2^2(\gamma_1-\gamma_2)^2)} = \frac{(1-\gamma_1)^2}{\sigma_3^2} \tag{S50}$$

$$\gamma_3 = \frac{\sigma_2^2(1-\gamma_1)^2(\gamma_1-\gamma_2)^2}{\sigma_3^2} \tag{S51}$$

$$\gamma_3 = \frac{S^{\text{center,surround}}(\sigma_\Delta^2 + C^{\text{group}}\sigma^2_{\text{prior}})^2}{[(\sigma_1^2\gamma_2 + C^{\,\text{surround}}\sigma^2_{\text{prior}} + \sigma_2^2(\gamma_1-\gamma_2)^2)]} \tag{S52}$$

$$\gamma_3 = -\frac{S^{\text{center,surround}}(\sigma_\Delta^2 + C^{\text{group}}\sigma^2_{\text{prior}})g}{[\sigma_\Delta^2 + \sigma_g^2 + S^{\text{center,surround}}(\sigma_\Delta^2 + C^{\text{group}}\sigma^2_{\text{prior}})]} \tag{S53}$$

$$\frac{(1-\gamma_1)}{(\gamma_1-\gamma_2)}\gamma_3 = -\frac{S^{\text{center,surround}}(\sigma_\Delta^2 + C^{\text{group}}\sigma^2_{\text{prior}})}{(\sigma_1^2\gamma_2 + C^{\,\text{surround}}\sigma^2_{\text{prior}} + \sigma_2^2(\gamma_1-\gamma_2)^2)} \tag{S54}$$

$$g = \frac{(1-\gamma_1)}{(\gamma_1-\gamma_2)}\gamma_3 \tag{S55}$$

$$g = -\frac{S^{\text{center,surround}}(\sigma_\Delta^2 + C^{\text{group}}\sigma_{\text{prior}}^2)}{\sigma_\Delta^2 + \sigma_r^2 + C^{\text{ surround}}\sigma_{\text{prior}}^2 + S^{\text{center,surround}}(\sigma_\Delta^2 + C^{\text{group}}\sigma_{\text{prior}}^2)} \tag{S56}$$

Substituting in the original expression

$$\begin{gathered} p(\vec{o}^{\text{ center}}, \vec{o}^{\text{ surround}}, \vec{v}^{\text{center}}_{\text{relative}} | C^{\text{ center}}, S^{\text{center,surround}}, C^{\text{group}}, C^{\text{ surround}}) = \\ (1-\gamma_1)^2 \mathcal{N}(\vec{o}^{\text{ surround}}(1-\gamma_1); 0, \sigma_3^2)\mathcal{N}(\vec{v}^{\text{center}}_{\text{relative}}; \vec{o}^{\text{ center}} + \vec{o}^{\text{ surround}} g, \sigma_2^2(1-\gamma_1)^2(1-S^{\text{center,surround}}\gamma_3)) \\ \mathcal{N}(\vec{v}^{\text{center}}_{\text{relative}}; 0, C^{\text{ center}}\sigma_{\text{prior}}^2) \end{gathered} \tag{S57}$$

By defining

$$\sigma_4^2 = \sigma_2^2(1-\gamma_1)^2(1-S^{\text{center,surround}}\gamma_3) + C^{\text{ center}}\sigma_{\text{prior}}^2 \tag{S58}$$

$$\gamma_4 = \frac{C^{\text{ center}}\sigma_{\text{prior}}^2}{\sigma_4^2} \tag{S59}$$

We can apply the rule for multiplying Gaussians and substitute the expression in the definition of the posterior of relative center velocity to get

$$\begin{gathered} p(\vec{v}^{\text{center}}_{\text{relative}} | \vec{o}^{\text{ center}}, \vec{o}^{\text{ surround}}, C^{\text{ center}}, S^{\text{center,surround}}, C^{\text{group}}, C^{\text{ surround}}) = \\ \mathcal{N}(\vec{v}^{\text{center}}_{\text{relative}}; (\vec{o}^{\text{ center}} + \vec{o}^{\text{ surround}} g)\gamma_4, \sigma_2^2(1-\gamma_1)^2(1-S^{\text{center,surround}}\gamma_3)\gamma_4) \end{gathered} \tag{S60}$$

which can be simplified as follows

$$\begin{gathered} p(\vec{v}^{\text{center}}_{\text{relative}} | \vec{o}^{\text{ center}}, \vec{o}^{\text{ surround}}, C^{\text{ center}}, S^{\text{center,surround}}, C^{\text{group}}, C^{\text{ surround}}) = \\ \mathcal{N}(\vec{v}^{\text{center}}_{\text{relative}}; (\vec{o}^{\text{ center}} + \vec{o}^{\text{ surround}} g)\gamma_4, [\sigma_4^2 - C^{\text{ center}}\sigma_{\text{prior}}^2]\gamma_4) \end{gathered} \tag{S61}$$

$$\begin{gathered} p(\vec{v}^{\text{center}}_{\text{relative}} | \vec{o}^{\text{ center}}, \vec{o}^{\text{ surround}}, C^{\text{ center}}, S^{\text{center,surround}}, C^{\text{group}}, C^{\text{ surround}}) = \\ \mathcal{N}(\vec{v}^{\text{center}}_{\text{relative}}; (\vec{o}^{\text{ center}} + \vec{o}^{\text{ surround}} g)\gamma_4, C^{\text{ center}}\sigma_{\text{prior}}^2(1-\gamma_4)) \end{gathered} \tag{S62}$$

#### D Distribution over responses under perceptual estimates

The distribution over responses can be evaluated for the different perceptual estimates by substituting the different estimates detailed in the main text (Section: Perceptual estimation) into Eq. 16.

**Posterior sampling**

$$p(R|\underline{\nu}) = \sum_j \sum_{i=0}^{N} w_j^{\text{quad}} w_i'(\vec{\underline{o}}) \int_\theta \mathcal{N}_{\text{circular}}(R; \theta + b, \kappa_m) \mathcal{N}_{\text{circular}}[\theta; \mu_i'(\vec{\underline{o}}), \kappa_i'] \tag{S63}$$

which can be simplified using the product rule of von-Mises pdf (Murray and Morgenstern, 2010) to get

$$p(R|\underline{\nu}) = \sum_j \sum_{i=0}^{N} w_j^{\text{quad}} w_i'(\vec{\underline{o}}) \frac{I_0[\sqrt{(\kappa_i')^2 + \kappa_m + 2\kappa_i'\kappa_m \cos(R - b - \mu_i'(\vec{\underline{o}}))}]}{2\pi I_0(\kappa_m) I_0(\kappa_i')} \tag{S64}$$

**Structure sampling**

$$p(R|\underline{\nu}) = \sum_j \sum_{i=0}^{N} w_j^{\text{quad}} w_i'(\vec{\underline{o}}) \mathcal{N}_{\text{circular}}(R; \mu_i'(\vec{\underline{o}}) + b, \kappa_m). \tag{S65}$$

**Model averaging**

$$p(R|\underline{\nu}) = \sum_j w_j^{\text{quad}}(\vec{\underline{o}}) \mathcal{N}_{\text{circular}} \left( R; \sum_{i=0}^{N} w_i' \mu_i'(\vec{\underline{o}}) + b, \kappa_m \right) \tag{S66}$$

**Model selection**

$$p(R|\underline{\nu}) = \sum_j w_j^{\text{quad}}(\vec{\underline{o}}) \mathcal{N}_{\text{circular}} \left( R; \mu_{i^*}'(\vec{\underline{o}}) + b, \kappa_m \right) \tag{S67}$$

### E Simplified model predictions for 3 moving elements

While having 3 different groups of dots results in 264 possible structures (Methods, Figure S9), for tractability and ease of interpretation we restricted the structures examined to those in which one of the relative velocities was zero. In addition, we assumed that the uncertainty in the inner ring's velocity was smaller than that of the center ring, that the uncertainty in the outer ring's velocity was smaller than that of the inner ring, and that all uncertainties were smaller than the uncertainty in the prior over relative velocities. Under these assumptions, the velocity of the group formed by any two elements is: (a) the velocity of the more reliable element if both are stationary relative to the group, and (b) the velocity of the stationary element if the other is moving. This simplification was possible due to the fact that any group of two moving elements in which one of them is stationary relative to the group, has the group velocity determined by the retinal velocity of the stationary element. The number of relevant structures under this simplification reduces to 44 principal structures.

This simplified model has 13 parameters which are: (a) the three sensory uncertainties associated with each velocity (3 parameters), (b) the computational noise at each level of the hierarchy in the model (1 parameter), (c) the mixture prior parameters (2 parameters), (d) the probability of grouping center & inner ring, inner & outer ring, and center & outer ring (3 parameters), (e) the probability of grouping the groups in (d) with the remaining element (3 parameters), (f) the probability of grouping center, inner and outer ring together into a single group (1 parameter).

To generate synthetic observers, we started with plausible parameters informed by the model fits to Experiment 1, and by choosing the inner and outer ring uncertainties and the computational noise to be much smaller, and the prior to be much larger, than the center ring uncertainty in accordance with simplifying assumptions. We also used prior knowledge of the stimulus spatial arrangement to choose a higher probability of the center being grouped with the inner ring (compared to the other combinations) and then with the outer ring. The values for the parameters are given in Supplementary Table S2.

### Supplementary Figures

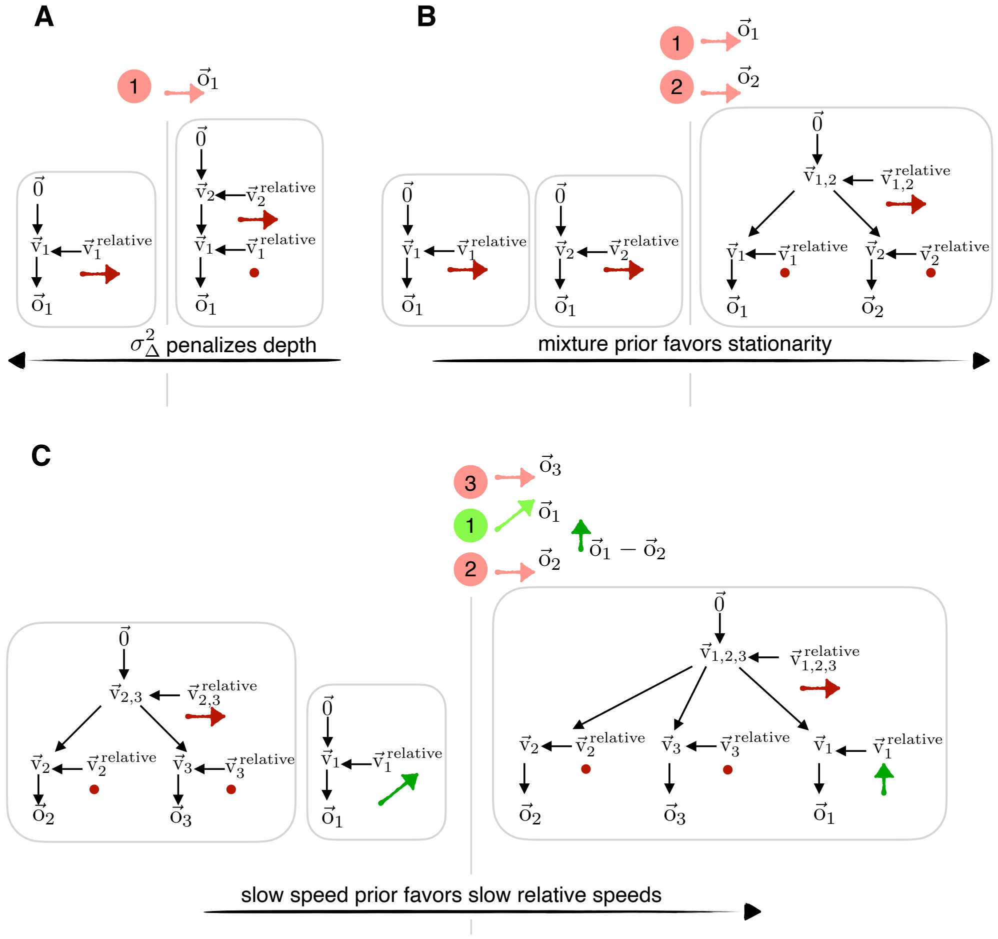


Figure S1: **Generative model in Figure 2 with the inferred velocities indicated.** Dot velocities and the corresponding inferred structures for stimuli with one moving dot (**A**), two moving dots (**B**) or three moving dots (**C**)

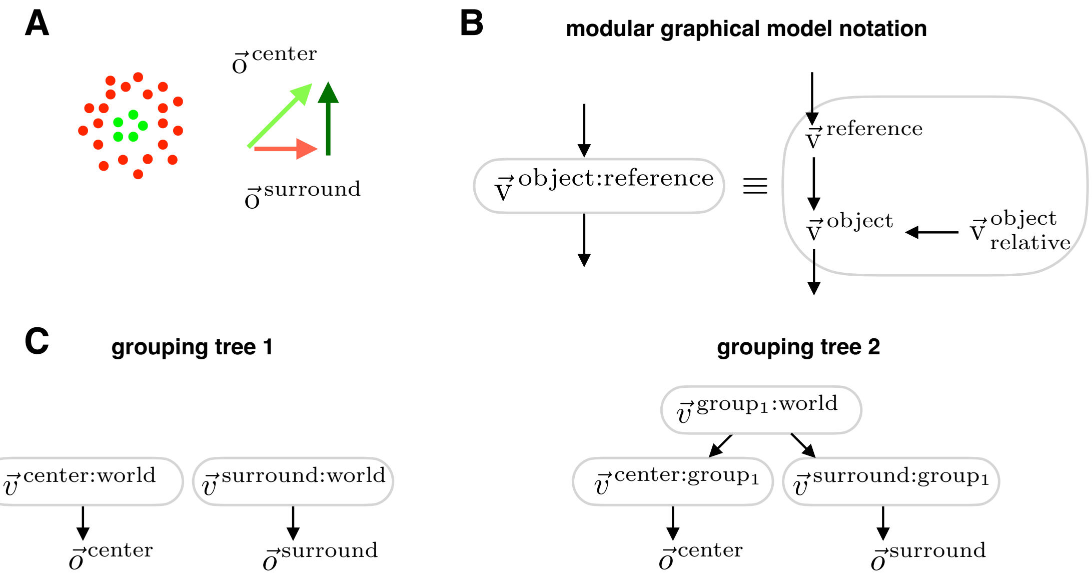


Figure S2: **Scenes with two moving elements.** **A** Stimulus with two coherently moving groups of dots whose velocities are chosen such that the target (green dots) moves perpendicularly to the surround (red dots). The stimulus is designed to separate three possible percepts of the target velocity as predicted by our model: (a) perceiving the retinal velocity (light green vector) if the target and group are not inferred to be part of the same motion structure (b) perceiving the group velocity (red vector) if the target is inferred to be part of the same motion structure as the surround and the target moves with the surround (c) perceiving the relative velocity (dark green vector) if the target is inferred to be part of the same motion structure as the surround but the target moves differently from the surround **(B)** A compact notation for the causal inference motif where an object is inferred in a reference frame by inferring whether its relative velocity to the reference frame is zero or not. **C** The different grouping trees that the model can infer with two moving elements.

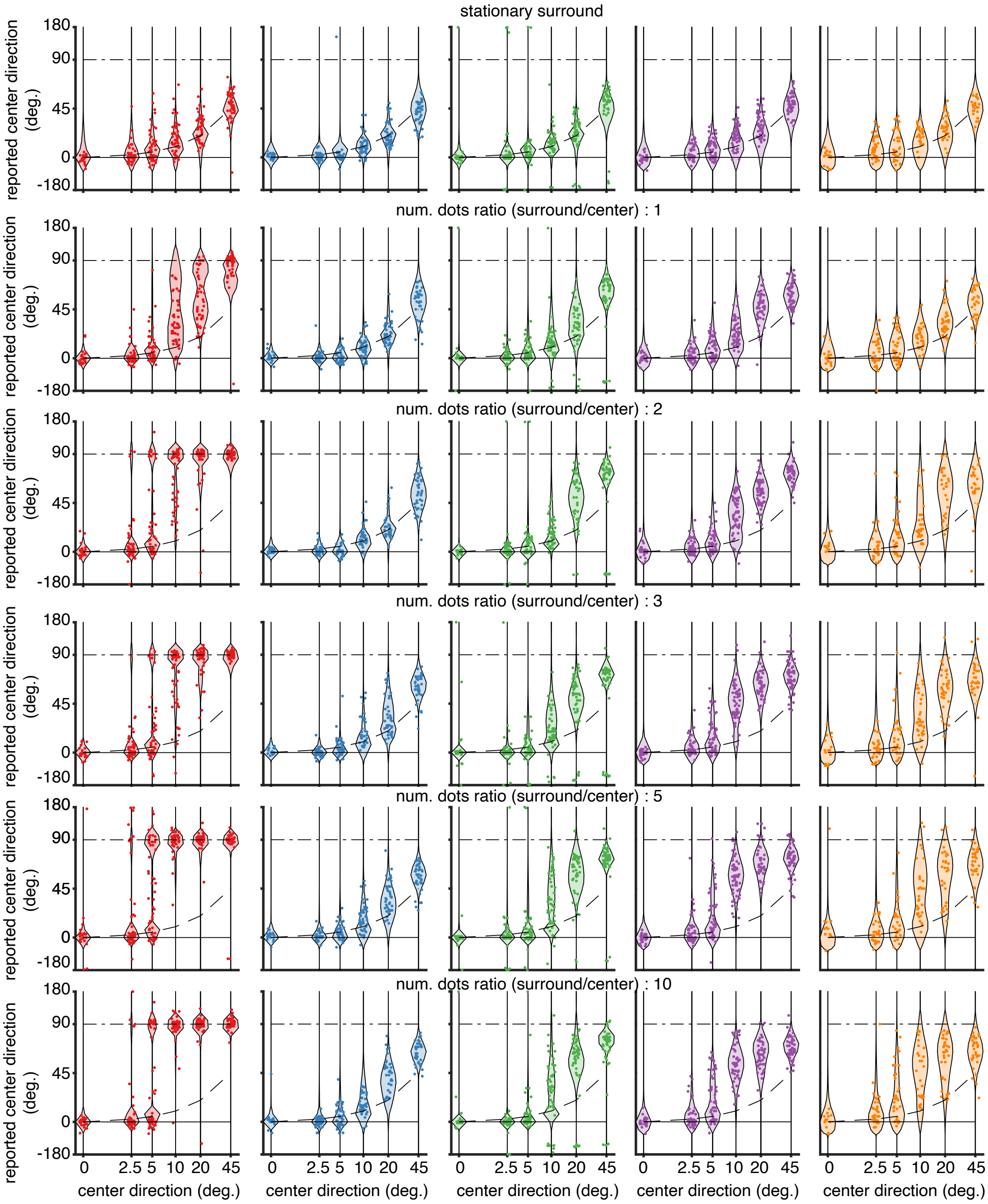


Figure S3: **Individual observer responses in experiment 1.** Response across all observers and different ratios of surround to center dots along with best fit model predictions. Each color indicates a different observer and each row corresponds to a different number of surround patches (indicated by the title).

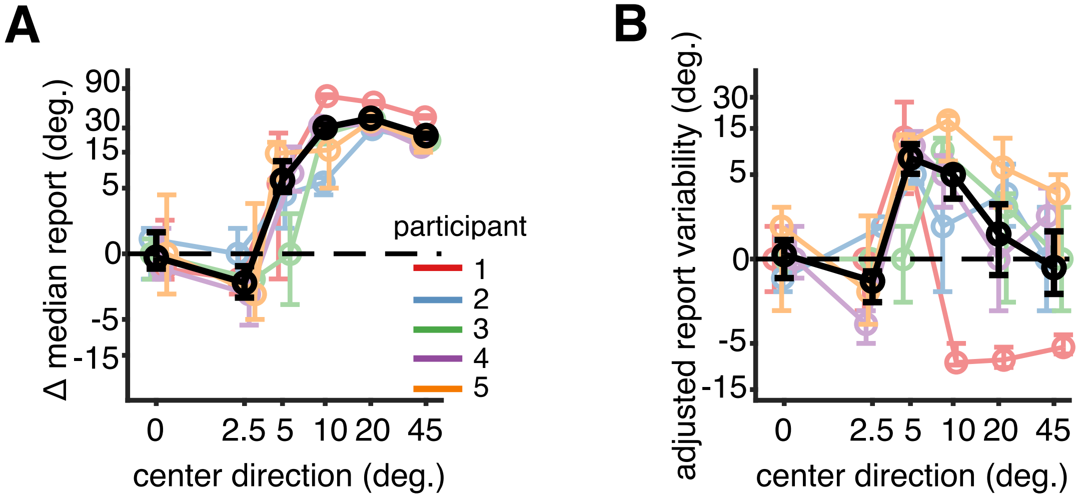


Figure S4: **Summary statistics of observer responses.** **(A)** Difference in median reports between moving and stationary surround conditions with 68% confidence intervals. Negative values indicate integration and positive values indicate segmentation.**(B)** Difference in report variability between moving and stationary surround with 68% confidence intervals, quantified by the difference in median absolute deviation (MAD). The MAD (Russell and Bernard, 2006) is a robust estimate of variability defined as the median absolute difference between individual trial reports and median reports across trials. Each color corresponds to a different observer and the black line shows the average across observers. The average difference in median reports is consistently negative for 2.5° (not significant with $p = 0.07$ across observers, with $p < 0.05$ individually for 3 out of 5 observers) indicating integration, and is significantly positive for larger separations (greater than 10 degrees; $p < 0.001$ across observers, also $p < 0.001$ individually for all observers) indicating segmentation. The small effect sizes for integration are a consequence of the experiment design as the maximum possible difference in median reports for a retinal center direction 2.5° is $-2.5°$ which is small compared to the reporting noise. This shortcoming is addressed in Experiment 2 described in the next section. The average MAD at 5° (Fig. 3D) is significantly greater than the MAD at 2.5° and 45° ($p < 0.001$ across observers, with $p < 0.05$ for 3 out of 5 observers individually) indicating greater variability in reports for intermediate separations reflecting the higher uncertainty in causal structures expected from the causal inference model.Significance was estimated by bootstrapping using $10^4$ samples.

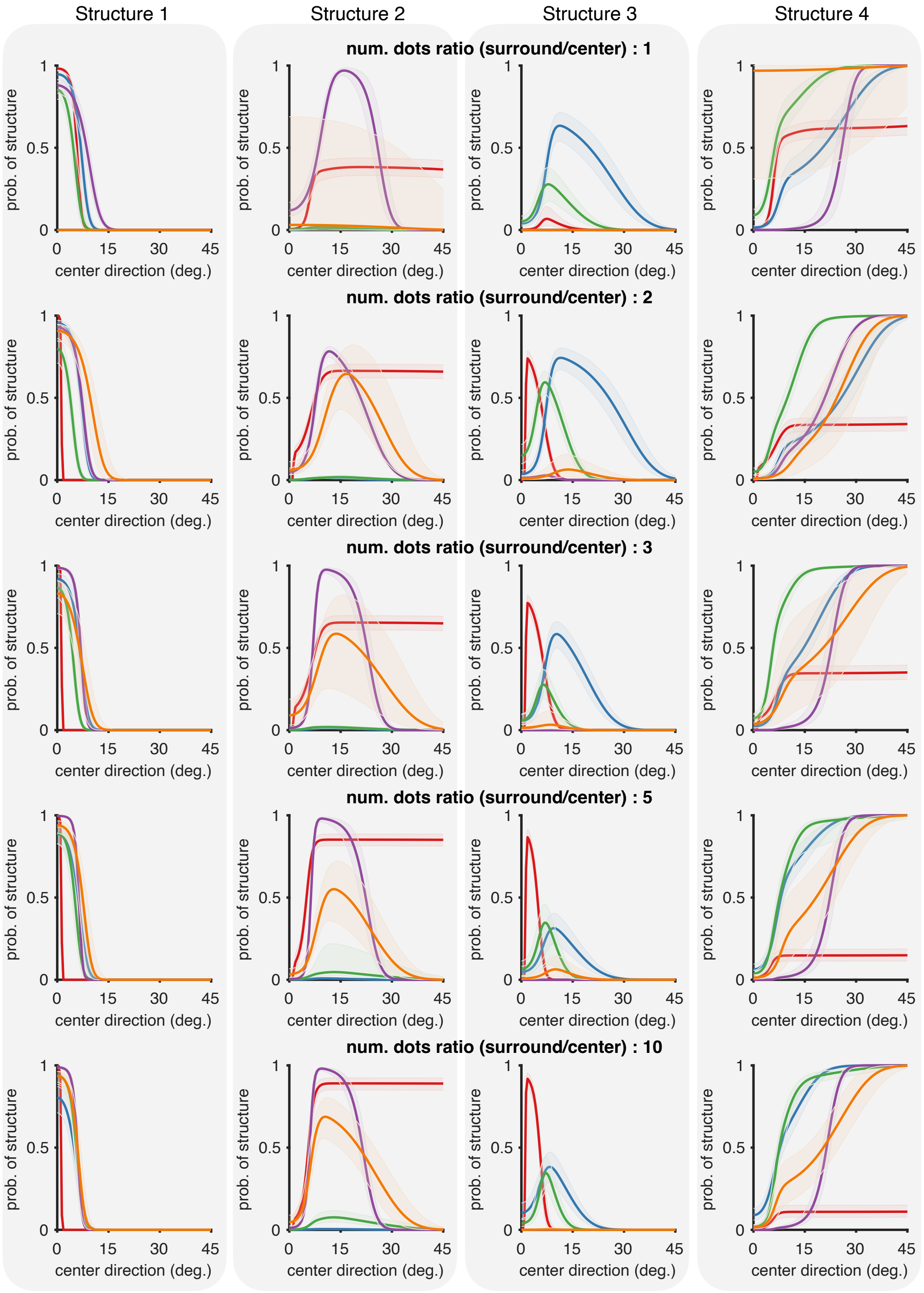

Structure 1
Structure 2
Structure 3
Structure 4
num. dots ratio (surround/center) : 1
num. dots ratio (surround/center) : 2
num. dots ratio (surround/center) : 3
num. dots ratio (surround/center) : 5
num. dots ratio (surround/center) : 10
prob. of structure
center direction (deg.)
0
0.5
1
15
30
45

Figure S5: **Posterior probability of structures for varying numbers of dots.** Posterior probability of the four structures in Figure 4A-D similar to those shown in Figure 4E-H for 10 surround patches, but for all numbers of surround patches (1,2,3,5 and 10).

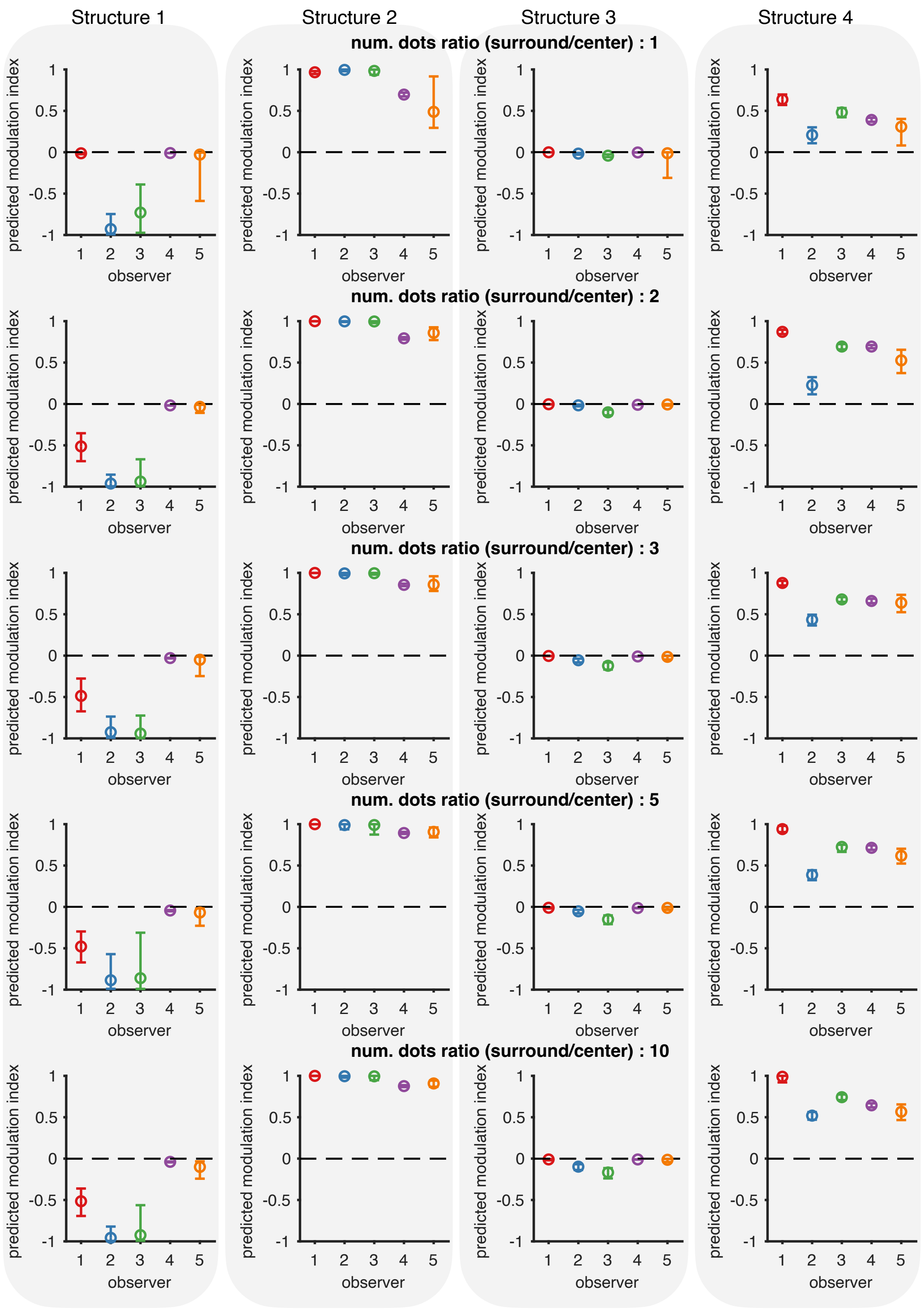
Structure 1
Structure 2
Structure 3
Structure 4
num. dots ratio (surround/center) : 1
num. dots ratio (surround/center) : 2
num. dots ratio (surround/center) : 3
num. dots ratio (surround/center) : 5
num. dots ratio (surround/center) : 10
predicted modulation index
observer

Figure S6: **Modulation indices corresponding to each structure for varying numbers of dots.** Modulation indices predicted under the four structures in Figure 4A-D similar to those shown in Figure 4I-L for 10 surround patches, but for all numbers of surround patches (1,2,3,5 and 10).

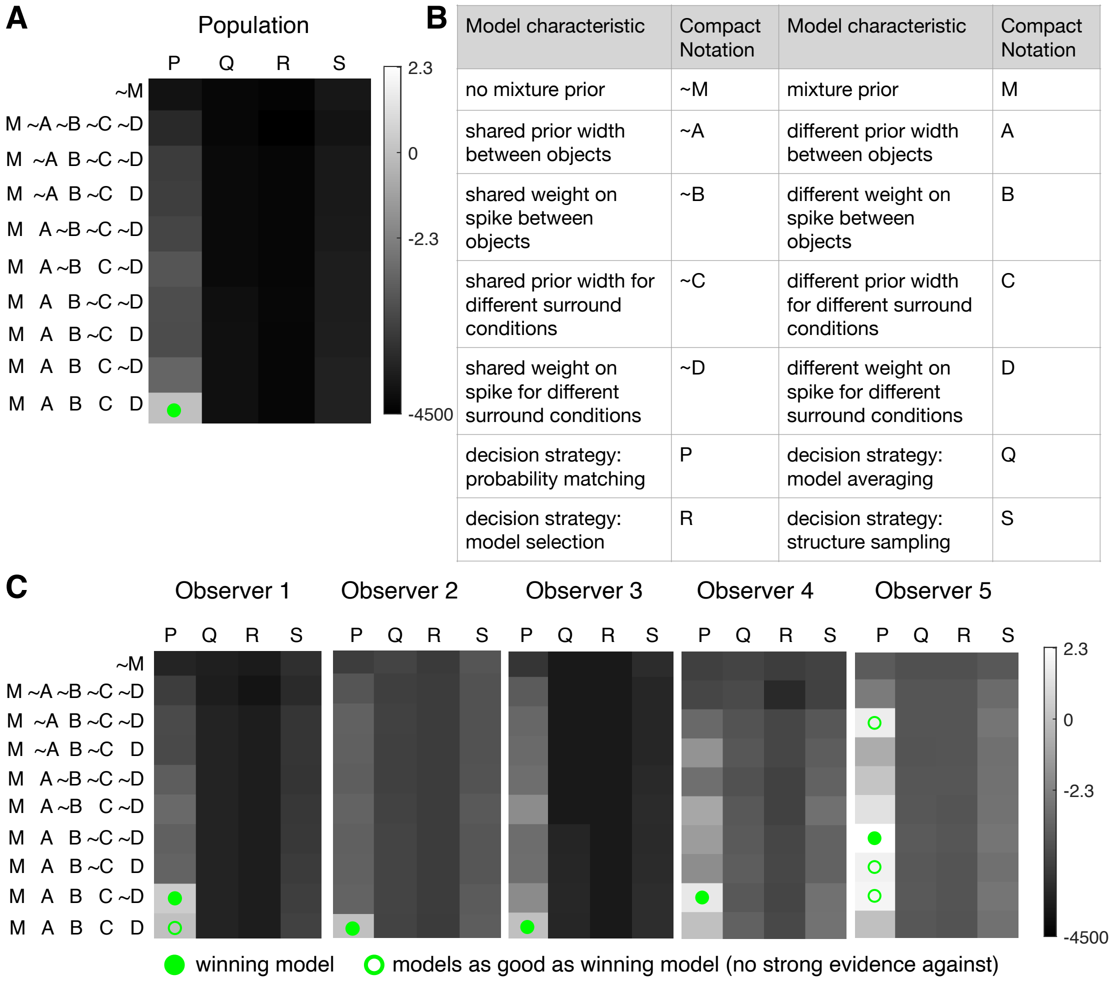


| Model characteristic | Compact Notation | Model characteristic | Compact Notation |
|---|---|---|---|
| no mixture prior | ~M | mixture prior | M |
| shared prior width between objects | ~A | different prior width between objects | A |
| shared weight on spike between objects | ~B | different weight on spike between objects | B |
| shared prior width for different surround conditions | ~C | different prior width for different surround conditions | C |
| shared weight on spike for different surround conditions | ~D | different weight on spike for different surround conditions | D |
| decision strategy: probability matching | P | decision strategy: model averaging | Q |
| decision strategy: model selection | R | decision strategy: structure sampling | S |

Figure S7: **Factorial model comparison.** Factorial model comparison using relative log likelihood (shown in grayscale) computed using AIC as in Figure 4B but across all simplifications of the model depicted in **(B)** shown for each observer in **(C)** and across all observers in **(A)**. The filled green circle indicates the best model through model comparison and the open green circles show models that are not the best but the evidence against them is not strong as compared to the best model. The most simplified model had the same prior parameters (weight on the delta and width of the slow speed prior) for: (a) the different velocities variables (i.e. center, surround, group) in our model, and (b) the inferred velocities for stimuli with different numbers of surround patches. We systematically allowed these parameters to vary to get to the most complex model where all parameters were allowed to vary (but with a weak prior preferring a shared value).

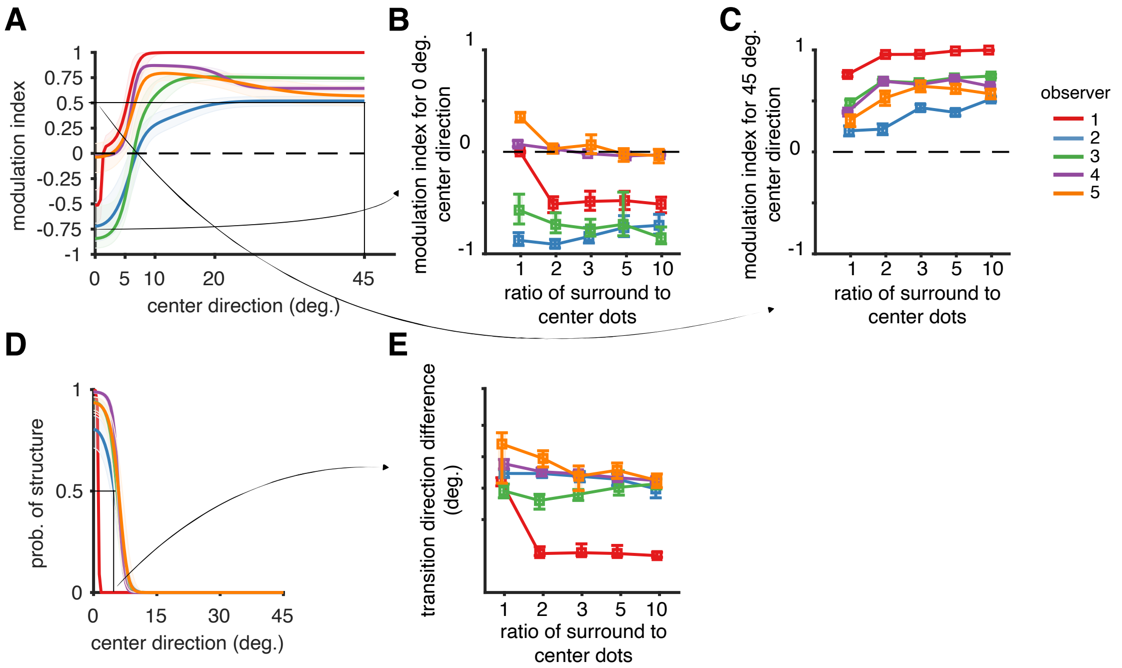


Figure S8: **Experiment 1: dependency on number of surround dots. (A)** The posterior predictive distribution over modulation indices across observers. **(B)** Predicted modulation index at 0° from (A) plotted as a function of the number of surround patches. 4/5 observers show a decrease in the predicted modulation index at 0 degrees (3 significantly with $p < 0.001$) with increasing number of surround patches indicating a higher strength of integration as the reliability of the surround increases. **(C)** Predicted modulation index at 45° from (A) plotted as a function of the number of surround patches. All observers show a significant increase in modulation index ($p < 0.001$) with increasing number of surround patches indicating a higher strength of segmentation as the reliability of the surround increases. The square markers in (B,C) indicate that these are model-predicted transition directions and not empirically measured data. All the statistical tests were done using samples obtained through posterior sampling of the model parameters. Two statistical tests were done to assess change in modulation indices as a function of the number of surround patches. First, the probability of modulation indices for 10 surround patches was estimated as being greater/lesser than those for 1 surround patch by comparing the posterior samples over the corresponding modulation indices. Second, the posterior slope distribution of the line fit to the modulation indices as a function of the number of dots was used to assess the increasing/decreasing trend by estimating the proportion of samples from the slope distribution that were greater/lesser than zero. **(D)** We quantify the region of integration by defining a transition direction difference which is the difference in direction between center and surround where the probability of integration is 0.5. The probability of integration is plotted in Figure S5 (first column) for different number of surround dots. **(E)** We find that the transition direction difference decreases with increase in number of surround dots in agreement with earlier causal inference studies that the region of integration increases with increase in cue uncertainty. The difference is significant ($p < 0.05$) for 4/5 observers.

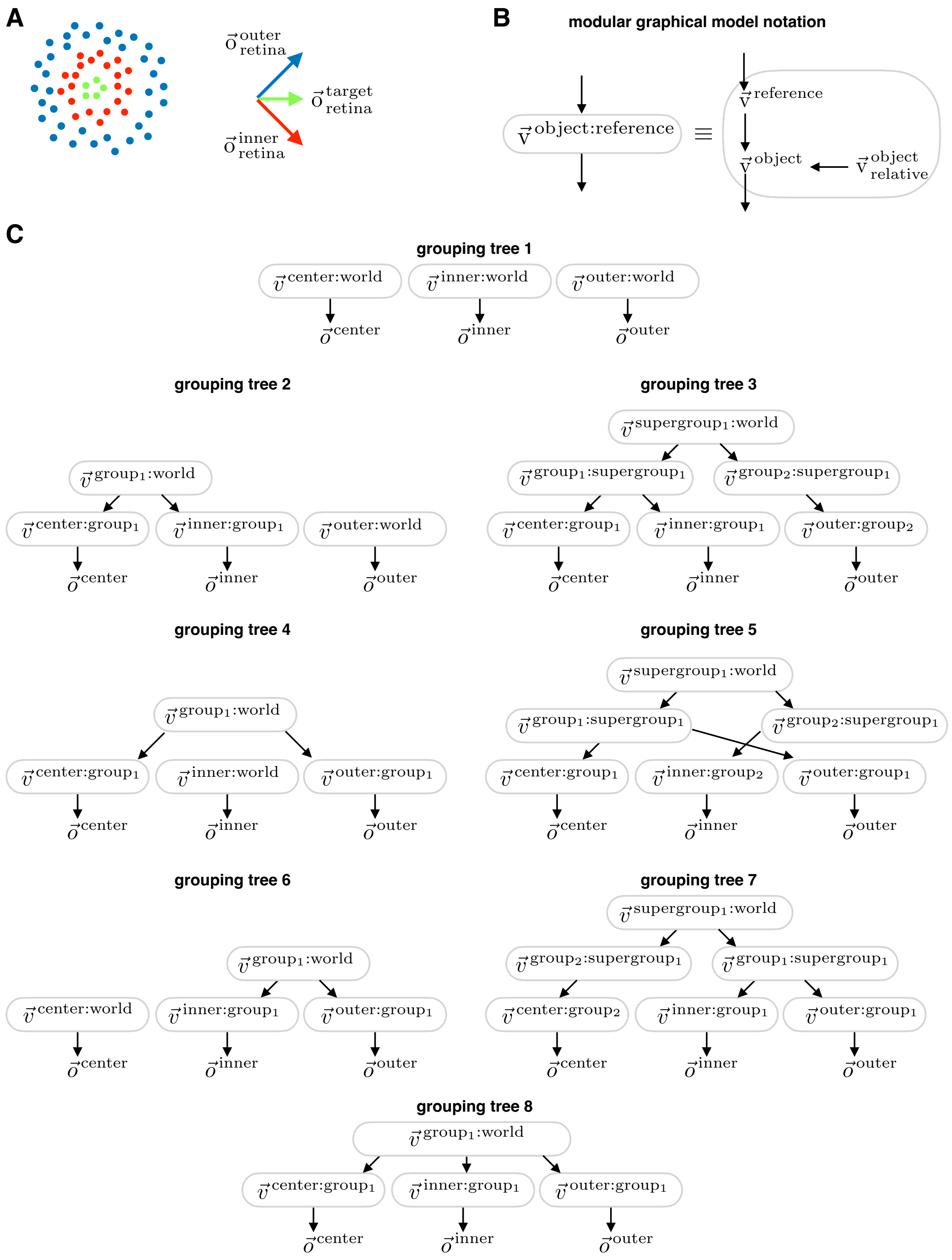
A
o outer retina
o target retina
o inner retina
B
modular graphical model notation
v object:reference
≡
v reference
v object
v object relative
C
grouping tree 1
v center:world
v inner:world
v outer:world
o center
o inner
o outer
grouping tree 2
v group1:world
v center:group1
v inner:group1
v outer:world
o center
o inner
o outer
grouping tree 3
v supergroup1:world
v group1:supergroup1
v group2:supergroup1
v center:group1
v inner:group1
v outer:group2
o center
o inner
o outer
grouping tree 4
v group1:world
v center:group1
v inner:world
v outer:group1
o center
o inner
o outer
grouping tree 5
v supergroup1:world
v group1:supergroup1
v group2:supergroup1
v center:group1
v inner:group2
v outer:group1
o center
o inner
o outer
grouping tree 6
v group1:world
v center:world
v inner:group1
v outer:group1
o center
o inner
o outer
grouping tree 7
v supergroup1:world
v group2:supergroup1
v group1:supergroup1
v center:group2
v inner:group1
v outer:group1
o center
o inner
o outer
grouping tree 8
v group1:world
v center:group1
v inner:group1
v outer:group1
o center
o inner
o outer

Figure S9: **Scenes with three moving elements.** **(A)** Stimulus with three coherently moving groups of dots whose velocities are such that the center (green dots) move perpendicularly to the surrounding inner and outer rings. The three velocities share a common component such that the relative velocities between any two elements are either 90 degrees or -90 degrees. **(B)** Replicated from Fig. S2B for clarity **(C)** The different causal structures that the model can infer with three moving elements. Each gray outline box implies two nested structures corresponding to whether the relative velocity is inferred to be zero or not. in **A**.

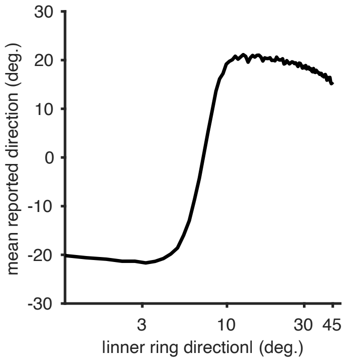


Figure S10: **Simplified model prediction for 3 moving elements.** We used the simplified model (Supplementary section E) to generate predictions for the mean responses (compare data in Figure 6B). The principle causal structures underlying this prediction, along with the predicted probability for each structure, are shown in Figures S11-S16.

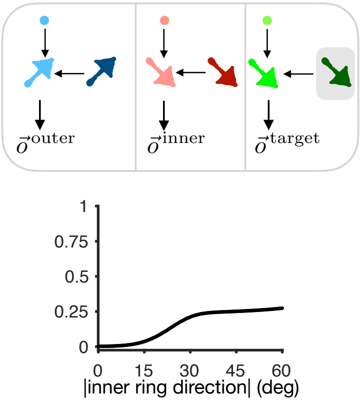


Figure S11: **Principle Structures for 3 Moving Elements: All elements moving independently.** See Figure S10 for initial context for the figure. In this figure the target (center), inner and outer rings move independently of each other.

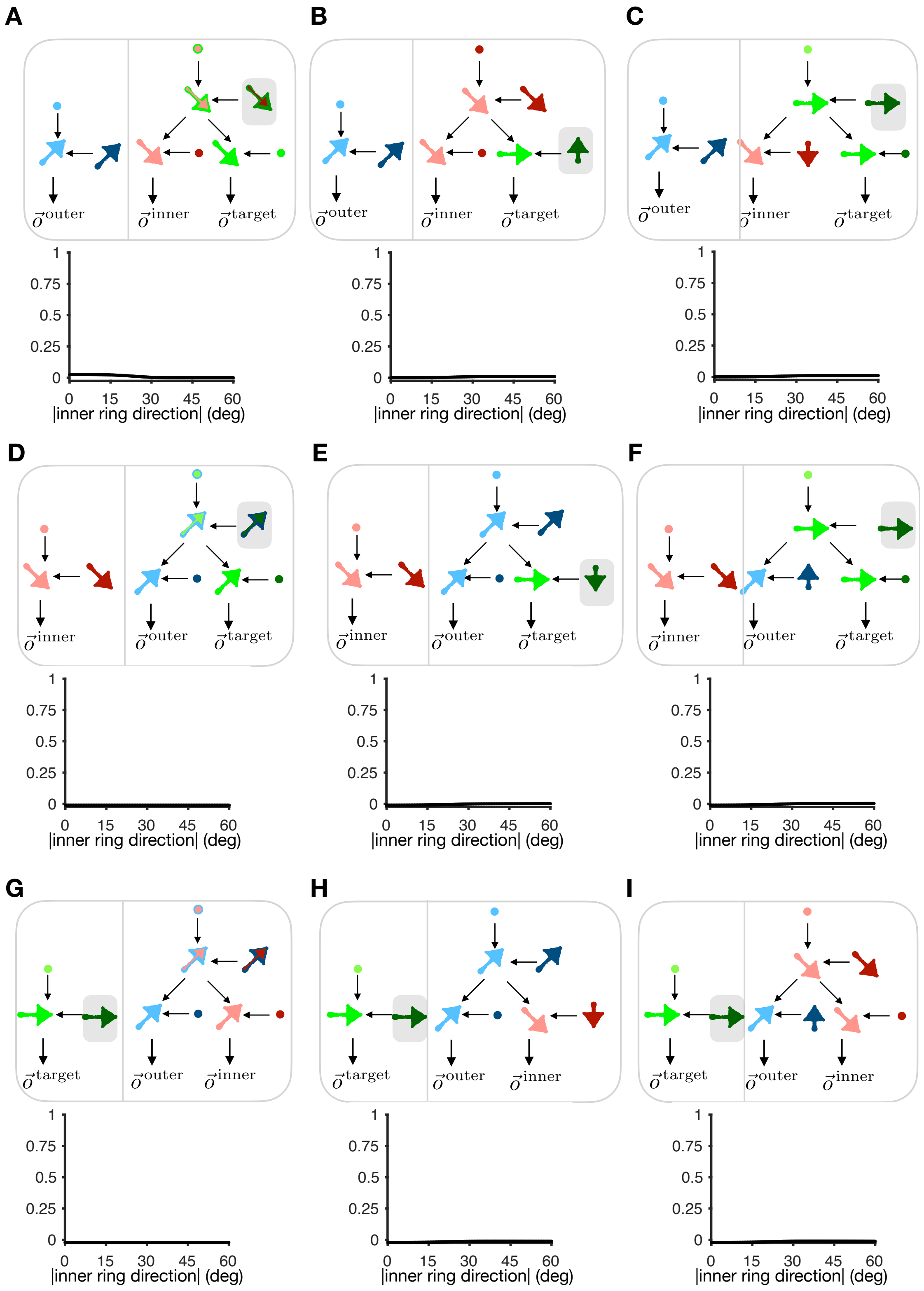
A
B
C
D
E
F
G
H
I
$\vec{o}^{outer}$
$\vec{o}^{inner}$
$\vec{o}^{target}$
1
0.75
0.5
0.25
0
0 15 30 45 60
|inner ring direction| (deg)

Figure S12: **Principle Structures for 3 Moving Elements: Two elements grouped together and other element moving independently** See Figure S10 for initial context for the figure. In this figure, two elements are grouped but the group is perceived independent of the third element. In the first row **(A-C)**, the center and inner rings form the group; in the second row **(D-F)**, the center and outer rings form the group; and in the third row **(G-I)** the inner and outer rings for the group

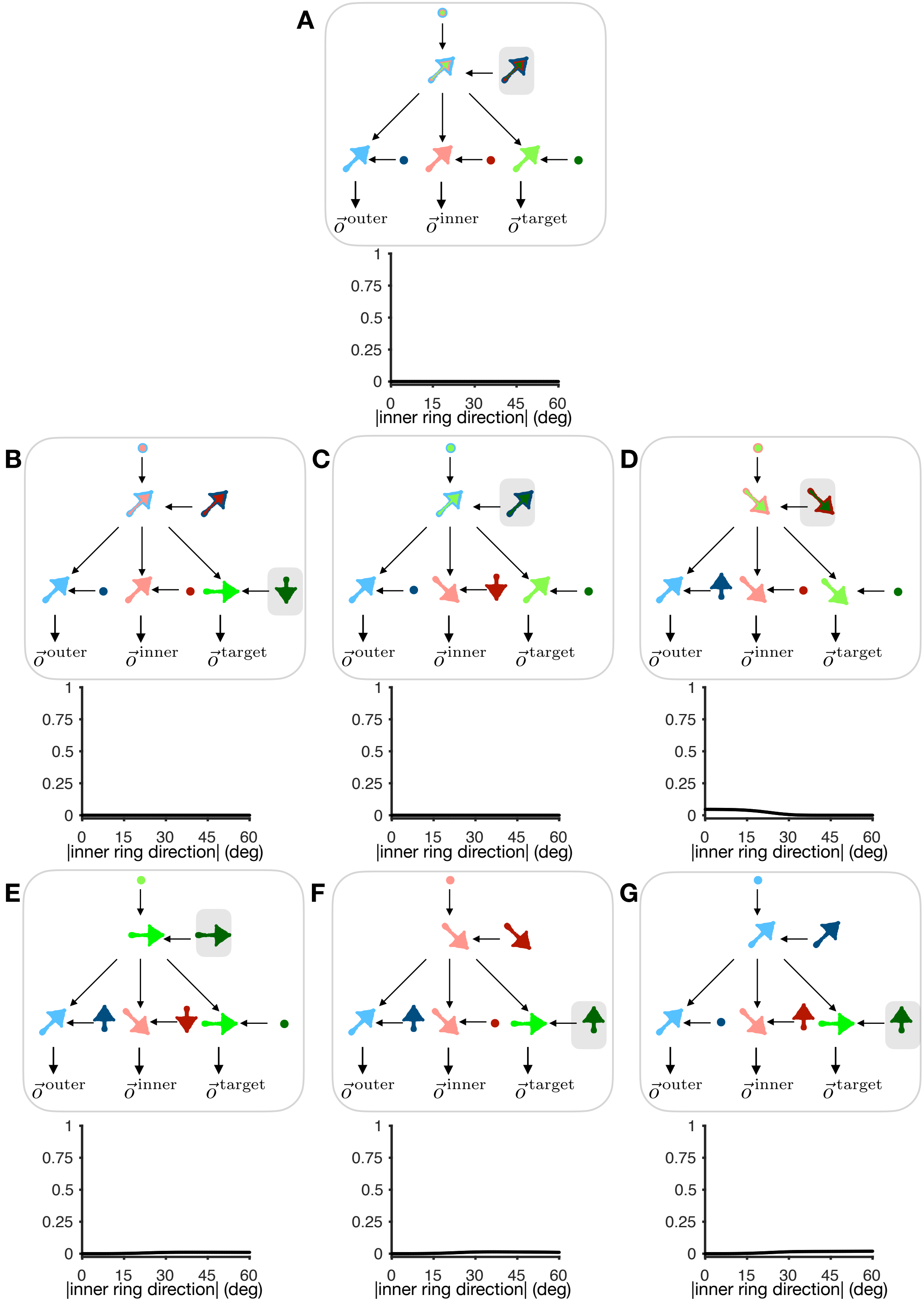

A
B
C
D
E
F
G
$\vec{o}^{\text{outer}}$
$\vec{o}^{\text{inner}}$
$\vec{o}^{\text{target}}$
1
0.75
0.5
0.25
0
0 15 30 45 60
|inner ring direction| (deg)

Figure S13: **Principle Structures for 3 Moving Elements: All elements grouped together.** See Figure S10 for initial context for the figure. In this figure, all three elements are grouped together into a common group.

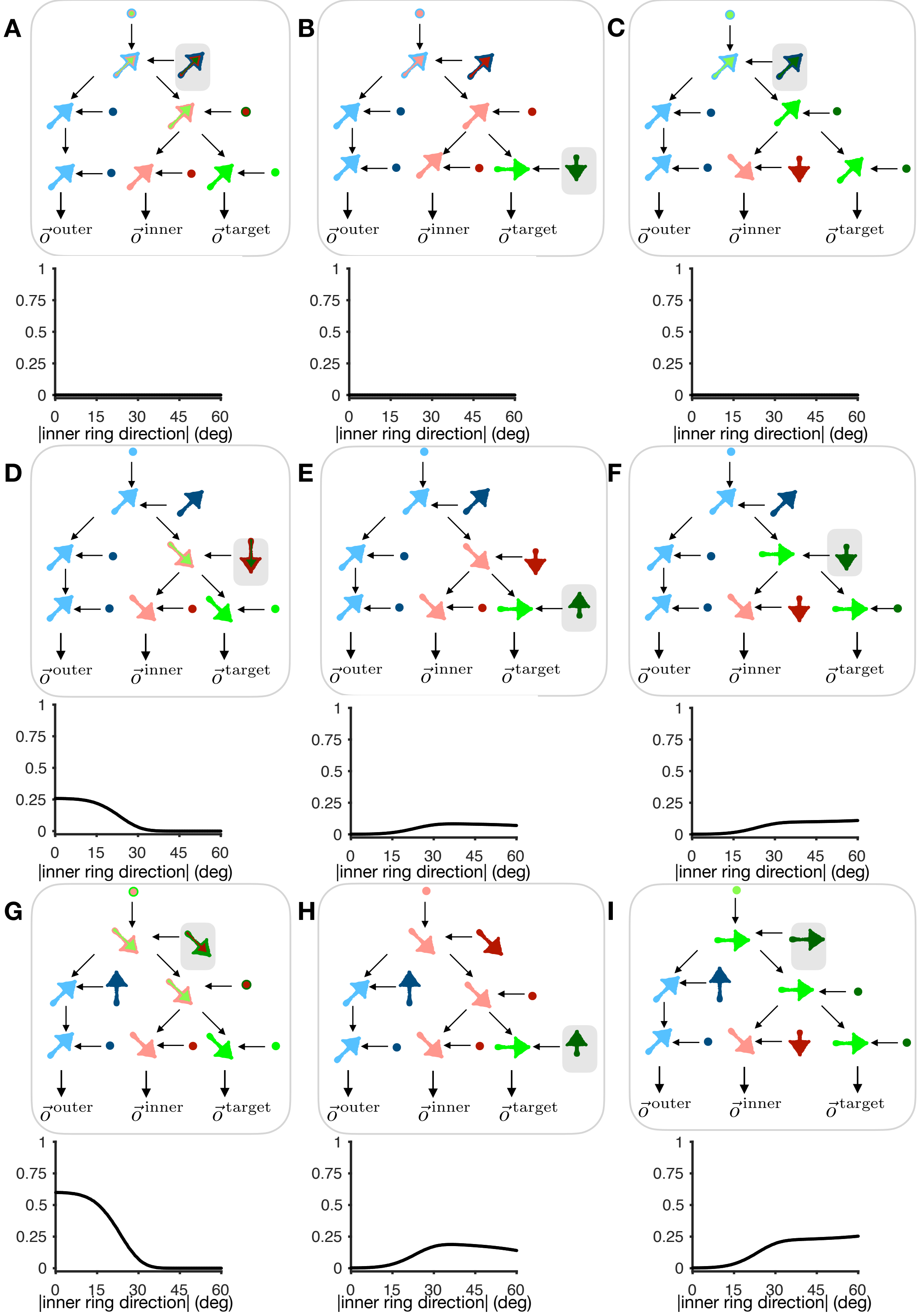
A
B
C
D
E
F
G
H
I
$\vec{o}^{outer}$
$\vec{o}^{inner}$
$\vec{o}^{target}$
0
0.25
0.5
0.75
1
0
15
30
45
60
|inner ring direction| (deg)

Figure S14: **Principle Structures for 3 Moving Elements: Two elements grouped together and the group forms a supergroup with the remaining element.** See Figure S10 for initial context for the figure. In this figure, center and inner rings are grouped and this group is grouped further with the outer ring forming a supergroup. Even though multiple structures have significant probability for the simulated parameters, only structures D and E make predictions for both center and surround velocities that match the responses in Figure S19. Future work fitting the model to data, will likely predict a higher probability for these two structures and a lower probability for the others.

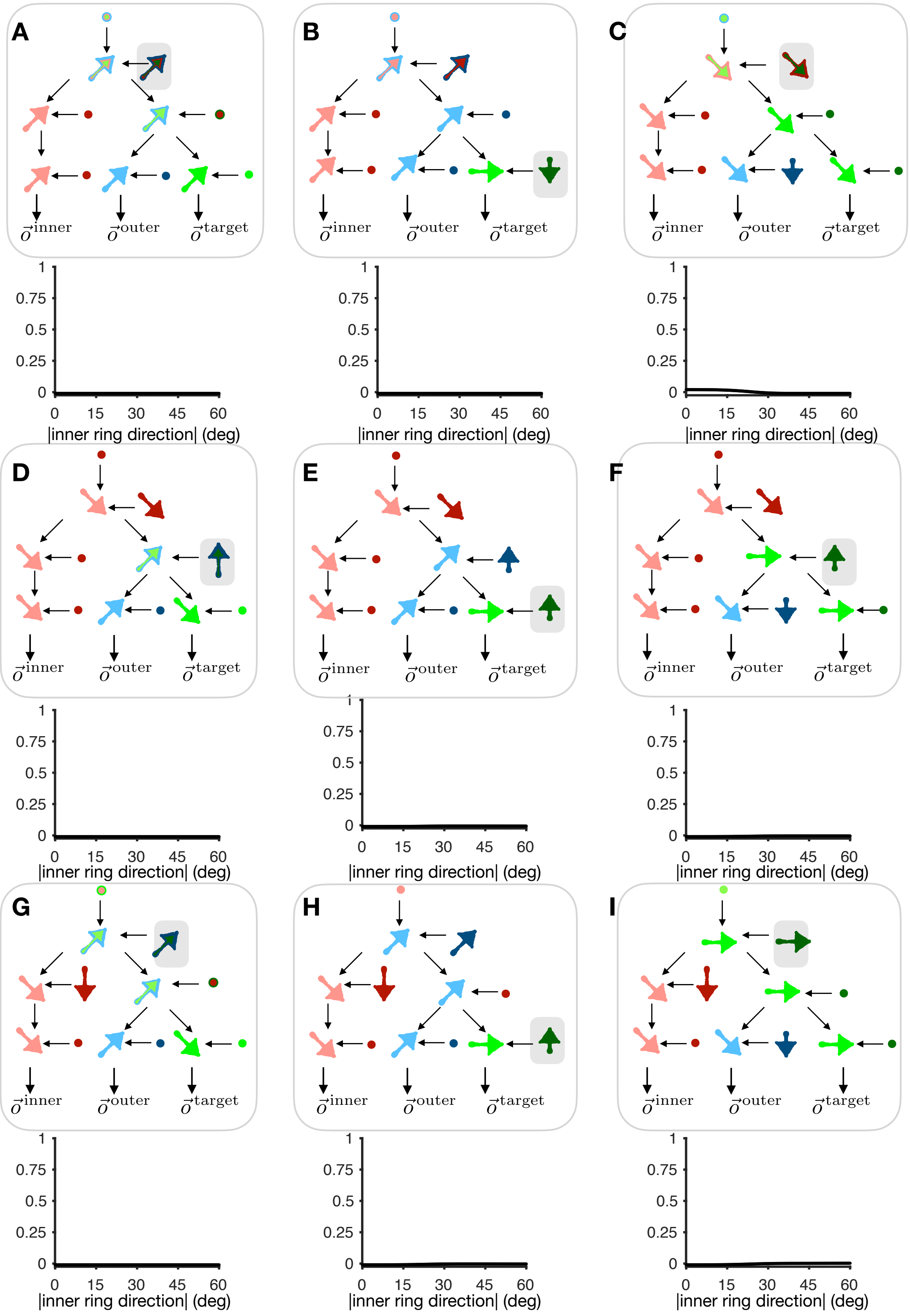

A
B
C
D
E
F
G
H
I
o⃗inner
o⃗outer
o⃗target
1
0.75
0.5
0.25
0
0 15 30 45 60
|inner ring direction| (deg)

Figure S15: **Principle Structures for 3 Moving Elements: Two elements grouped together and the group forms a supergroup with the remaining element.** See Figure S10 for initial context for the figure. In this figure, center and outer rings are grouped and this group is grouped further with the inner ring forming a supergroup.

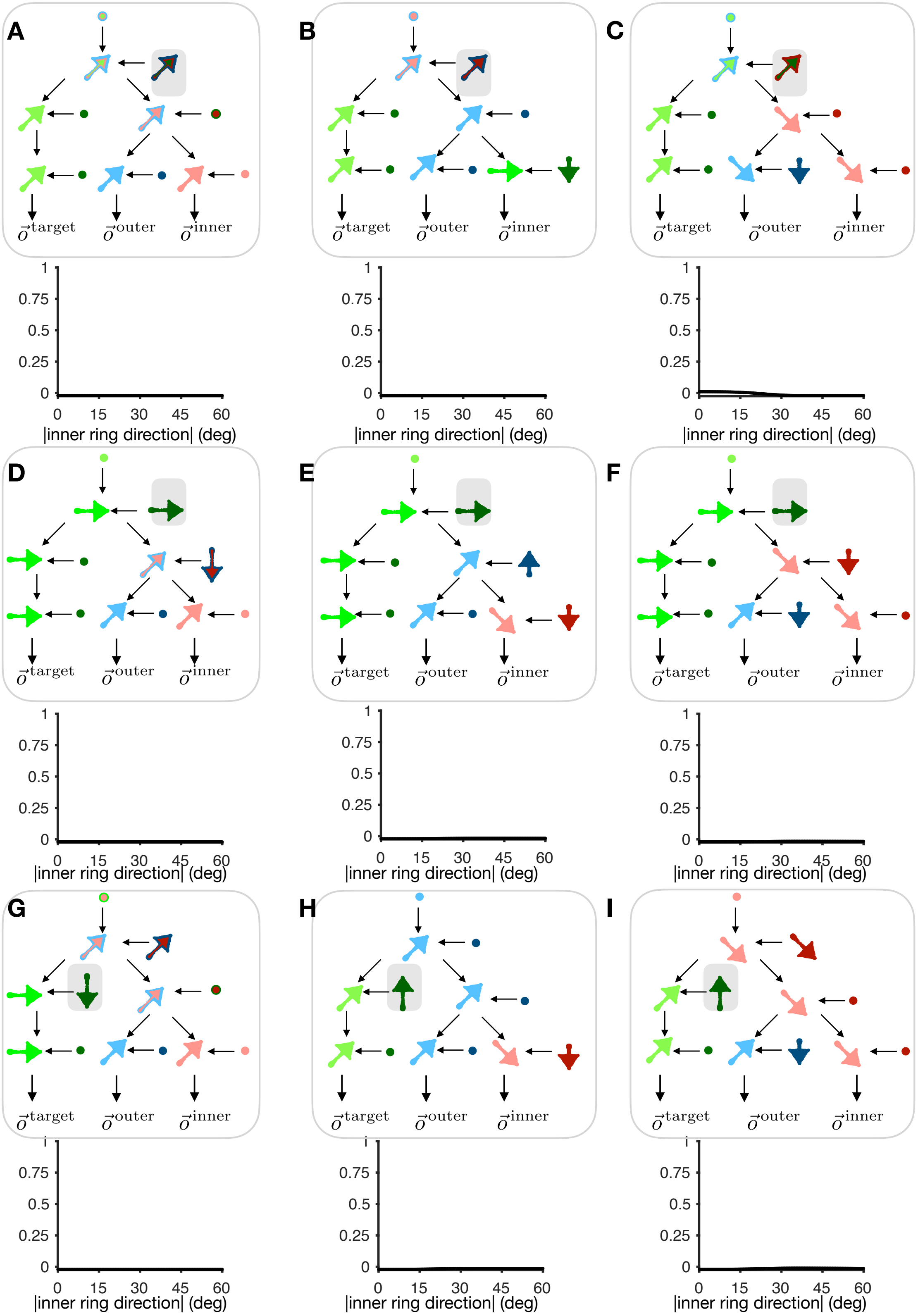
A
B
C
D
E
F
G
H
I
$\vec{o}^{\text{target}}$
$\vec{o}^{\text{outer}}$
$\vec{o}^{\text{inner}}$
1
0.75
0.5
0.25
0
0
15
30
45
60
|inner ring direction| (deg)

Figure S16: **Principle Structures for 3 Moving Elements: Two elements grouped together and the group forms a supergroup with the remaining element.** See Figure reffig:S-meanpred-three for initial context for the figure. In this figure, inner and outer rings are grouped and this group is grouped further with the center forming a supergroup.

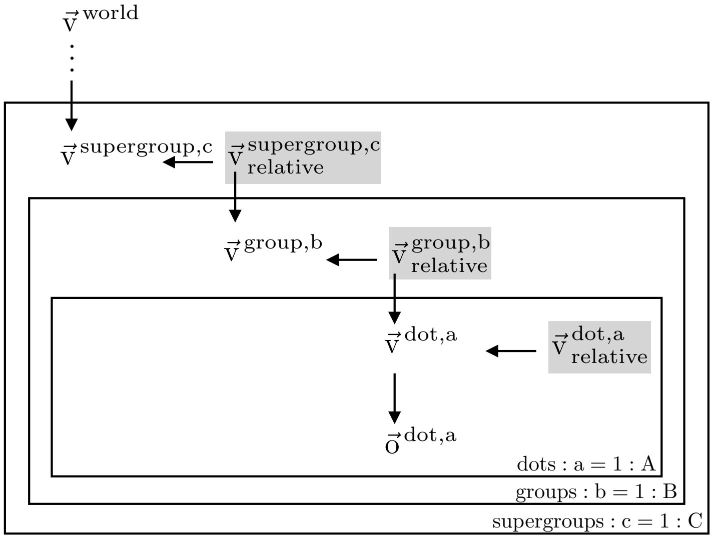


Figure S17: **Alternative (hypothetical) model for hierarchical motion perception.** The motif presented in Figure 1F could be stacked alternatively to Figure 1G such that the perceived velocity (gray shaded box) forms the reference frame for the level below. Such a model would be consistent with previous work in orientation perception (Chopin et al., 2012; Cicchini et al., 2021) where the perceived context value influences the perception of target stimuli. Experiment 1 cannot separate between this structure and Figure 1G as both the perceived and retinal group velocities are the same. But results from experiment 2 (with an additional level of hierarchy) flalsify this model and support our causal inference model presented in Figure 1G.

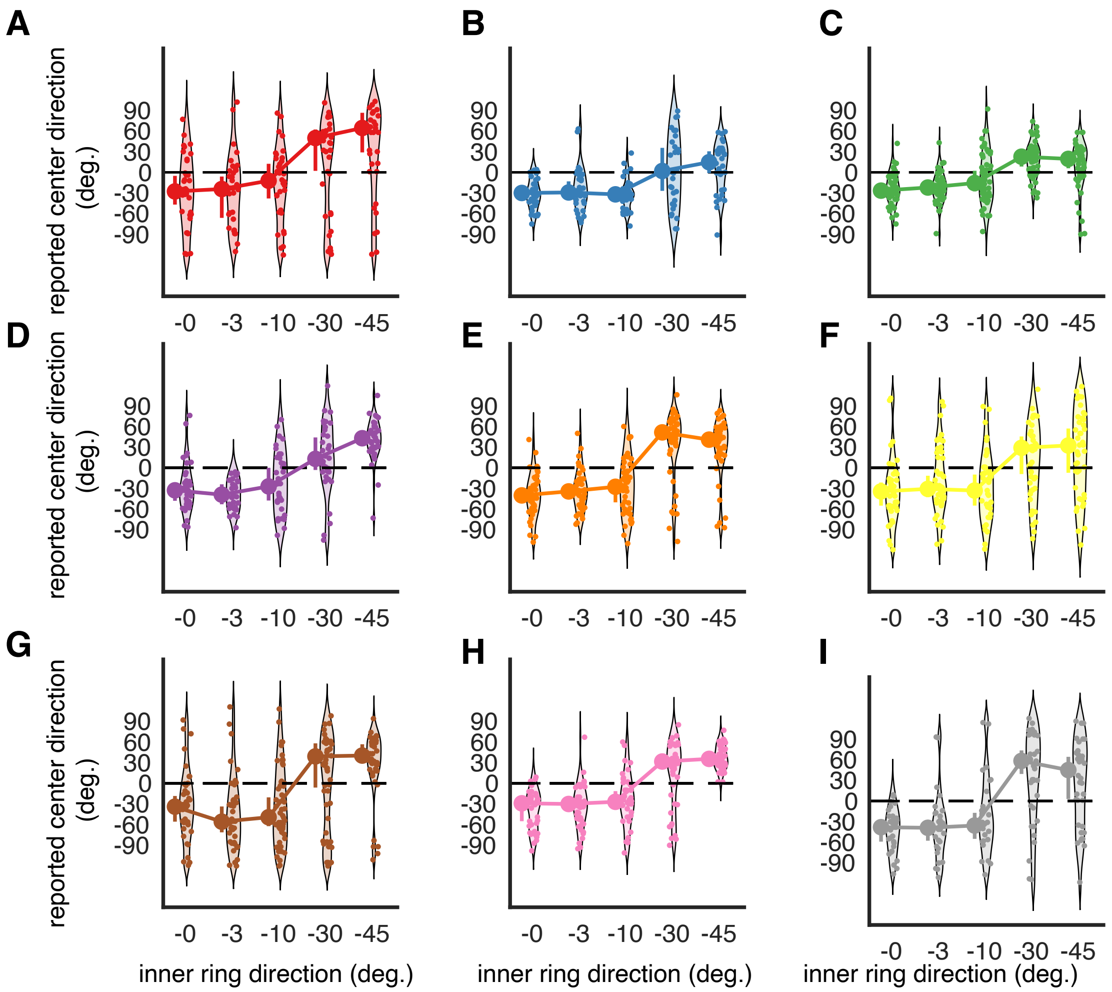


Figure S18: **Individual responses in experiment 2.** Individual responses across trials for observers in experiment 2 for the experimental condition shown in Figure 6A. Each dot corresponds to the response on a single trial and the violin plot is the empirical histogram of responses. The responses are summarized using the circular marginal median along with 95% CI as indicated by the errorbars to the left of each violin plot.

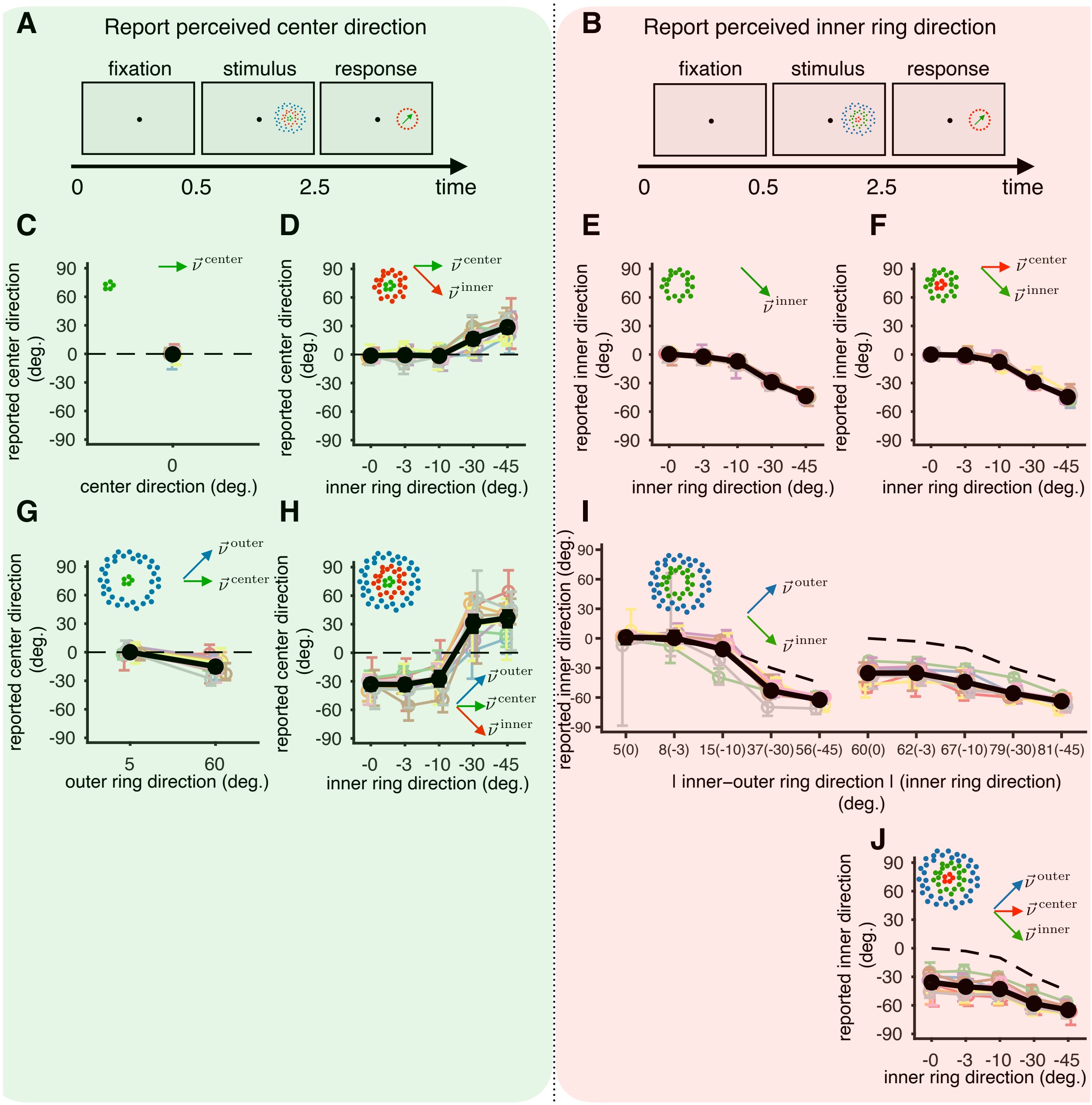
A
Report perceived center direction
fixation
stimulus
response
0
0.5
2.5
time
B
Report perceived inner ring direction
C
reported center direction (deg.)
center direction (deg.)
D
inner ring direction (deg.)
E
reported inner direction (deg.)
F
G
outer ring direction (deg.)
H
I
| inner–outer ring direction | (inner ring direction) (deg.)
J
-0 -3 -10 -30 -45
5(0) 8(-3) 15(-10) 37(-30) 56(-45) 60(0) 62(-3) 67(-10) 79(-30) 81(-45)
90 60 30 0 -30 -60 -90

Figure S19: **Additional empirical results with three moving elements. (A)** Stimulus schematic. The observer performs an estimation task in which they have to report the direction of green dots (center) using a dial. **(B)** Stimulus schematic. The observer performs an estimation task in which they have to report the direction of green dots (inner ring) using a dial.**(C,D,G,H)** Observer responses (and 95 % CI) for center percept for different stimulus conditions that define the velocities of the center, inner ring, and outer ring. The group of dots that are displayed in the inset are the coherently moving dots in that condition with velocities indicated by the vectors. The dots not displayed move randomly. The velocities are chosen such that the center always moves at 0 degrees (horizontally) and the two rings move with the same horizontal component but opposite vertical components such that the relative velocities are either 90 degrees (center and inner ring) or -90 degrees (center and outer ring). This arrangement is rotated around the circle across trials to account for any reporting biases. **(E,F,I,J)** Observer responses (and 95 % CI) for inner ring percepts for different stimulus conditions similar to **C,D,G,H**. The observer always reports the direction of the green dots and in these trials the green dots form the inner ring whereas they formed the center in **C,D,G,H**. For (I), the ticklabels indicate the absolute angular difference between the inner and outer ring directions and the brackets indicate the inner ring direction.
In two control conditions (C,E), the observers reported either the center or the inner ring's direction respectively, which was found to be aligned with the presented retinal direction demonstrating no reporting biases. When the center and inner ring moved coherently (D,F), the center direction reports followed the same pattern as predicted in experiment 1 (integration for small inner ring directions and segmentation for larger inner ring directions). The inner ring percepts (F) were largely unaffected by the center suggesting that the inner ring predominantly determines the group velocity. When the center and outer ring moved coherently (G), seven out of nine observers displayed significant negative biases. This outcome supports the notion of perceiving the relative velocity of the center to the outer ring, albeit with a low modulation index suggesting the effect of a surround on the center diminishes with increasing spatial separation between them. Similarly, when the inner and outer rings moved together (I), a weak integration was visible (flattening of the curve beyond the dashed line) for an absolute angular difference is 8 degrees (significant with $p<0.05$ across observers; significant with $p<0.05$ for 5/9 observers). We also see significant biases ($p<0.001$ across observers and also for each observer) consistent with perceiving the inner ring relative to the outer ring for larger separations. Reporting the inner ring directions in the condition in which all the elements move coherently (J), the observer responses are significantly negative across and for individual observers ($p<0.001$) consistent with the model predicts that the observers perceive the inner ring relative to the outer ring. The absolute angular differences between the inner and outer ring lie between 60 to 81 degrees and are similar to the responses when only the inner and outer rings move coherently (for large separations) supporting the model prediction that the percept of the inner ring is the perceived velocity of the center-inner group in the outer ring reference frame and that the group velocity is predominantly determined by the inner ring.

| Parameter | Prior | Remarks |
|---|---|---|
| $\sigma^2_{\text{prior}}$ | Lognormal$(-2, 2)$ | $\sim 90\%$ mass $\in [0.5°, 75°]$; |
| $\sigma^2_{\Delta}$ | Lognormal$(-2, 2)$ | see previous |
| $\sigma^2_{\text{center}}/\sigma^2_{\text{prior}}$ | Beta prime$(1, 1)$ | chosen this way such that weight on the observation in the posterior given by $\left(\frac{\sigma^2_{\text{prior}}}{\sigma^2_{\text{center}}+\sigma^2_{\text{prior}}}\right)$ is uniformly distributed |
| $\sigma^2_{\text{surround}}/\sigma^2_{\text{prior}}$ | Beta prime$(1, 1)$ | see previous |
| $\alpha$ | Beta$(1.25, 1.25)$ | weakly informative prior with a mode at 0.5 |
| $\beta^{\text{center,surround}}$ | Beta$(1.25, 1.25)$ | see previous |
| $\lambda$ | Beta$(1, 5)$ | prior incorporating knowledge that lapse rates are likely to be small |
| $\mu^{\text{lapse}}$ | Normal$(0, \pi)$ | symmetric prior over lapse mean probabilities in radians |
| $\kappa^{\text{lapse}}$ | Lognormal$(0, 2.35)$ | 95% mass $\in [0.01, 100]$ |
| $b$ | Normal$(0, \pi/2)$ | symmetric prior over response biasin radians |
| $\kappa_{\text{m}}$ | Lognormal$(0, 2.35)$ | 95% mass $\in [0.01, 100]$ |

Table S1: The weakly informative priors used over parameters for MAP and posterior estimation. When different parameters for $\sigma^2_{\text{prior}}$ are estimated for center and surround, and for different number of surround patches, a Normal prior $[\mathcal{N}(0, 4)]$ is placed on the difference between the parameters incorporating knowledge that apriori all parameters are likely to be similar for reduced displays. Similarly, a Normal prior $[\mathcal{N}(0, 0.5)]$ is placed on the log odds between the estimated $\alpha$ values for center and surround, and for different number of surround patches preferring similar estimates

| Parameter | Value for simulation |
|---|---|
| Target observation noise variance ($\sigma^2_{\text{target}}$) | $7.5 \times 10^{-3}$ |
| Inner ring observation noise variance ($\sigma^2_{\text{inner}}$) | $7.5 \times 10^{-5}$ |
| Outer ring observation noise variance ($\sigma^2_{\text{outer}}$) | $7.5 \times 10^{-11}$ |
| Variance of Gaussian component of the prior ($\sigma^2_{\text{prior}}$) | 10 |
| Computational noise variance ($\sigma^2_{\Delta}$) | $10^{-6}$ |
| Weight on delta component of prior ($\alpha$) | 0.7 |
| Prior probability over grouping target and inner ring ($\beta^{\text{target,inner}}$) | 0.8 |
| Prior probability over grouping target and outer ring ($\beta^{\text{target,outer}}$) | 0.05 |
| Prior probability over grouping inner and outer rings ($\beta^{\text{inner,outer}}$) | 0.05 |
| Prior probability over grouping target and inner ring group (group 1) with the outer ring ($\beta^{1,\text{outer}}$) | 0.99 |
| Prior probability over grouping target and outer ring group (group 2) with the inner ring ($\beta^{2,\text{inner}}$) | 0.8 |
| Prior probability over grouping inner and outer ring group (group 1) with the center ($\beta^{3,\text{center}}$) | 0.8 |
| Prior probability over grouping target, inner and outer into a single group ($\beta^{\text{all}}$) | 0.05 |

Table S2: Parameters used to generate simplified model predictions for experiment with 3 moving elements
